# NeuroEnergetics-on-Chip: a novel 3-compartment microfluidic platform to study metabolic interactions between brain parenchyma and cerebral vasculature

**DOI:** 10.64898/2026.06.21.733618

**Authors:** Angela Ceballos, Ludovica Montesi, Liisa Loel, Monika Yanovska, Justina Venckute, Jessika Jessika, Tingting Wu, Laura Benito-Zarza, John Cognetti, Omid Fotouhi, Kristaps Klavins, Sofia Ygberg, Anna Wredenberg, Anna Wedell, Anna Herland, Julia Rogal

## Abstract

Neurological disorders are a major cause of death and disability worldwide. The brain’s energy metabolism is essential to its proper function, yet the mechanisms driving neuroenergetic dysfunction remain poorly understood. A key challenge is the limited availability of human-relevant models that can reproduce the complexity of brain physiology. An Organ-on-Chip (OoC) system was developed to mimic the neurovascular unit metabolic coupling by incorporating human isogenic iPSC-derived endothelial-like cells, pericyte-like cells, astrocytes, and a cerebral organoid, representing the main cellular components of the NVU. The novel, customized microfluidic platform enables research on neurovascular coupling by interconnecting a blood-brain barrier-on-a-chip model with a 3D brain parenchymal compartment to mimic physiological conditions.

**Graphical abstract:** 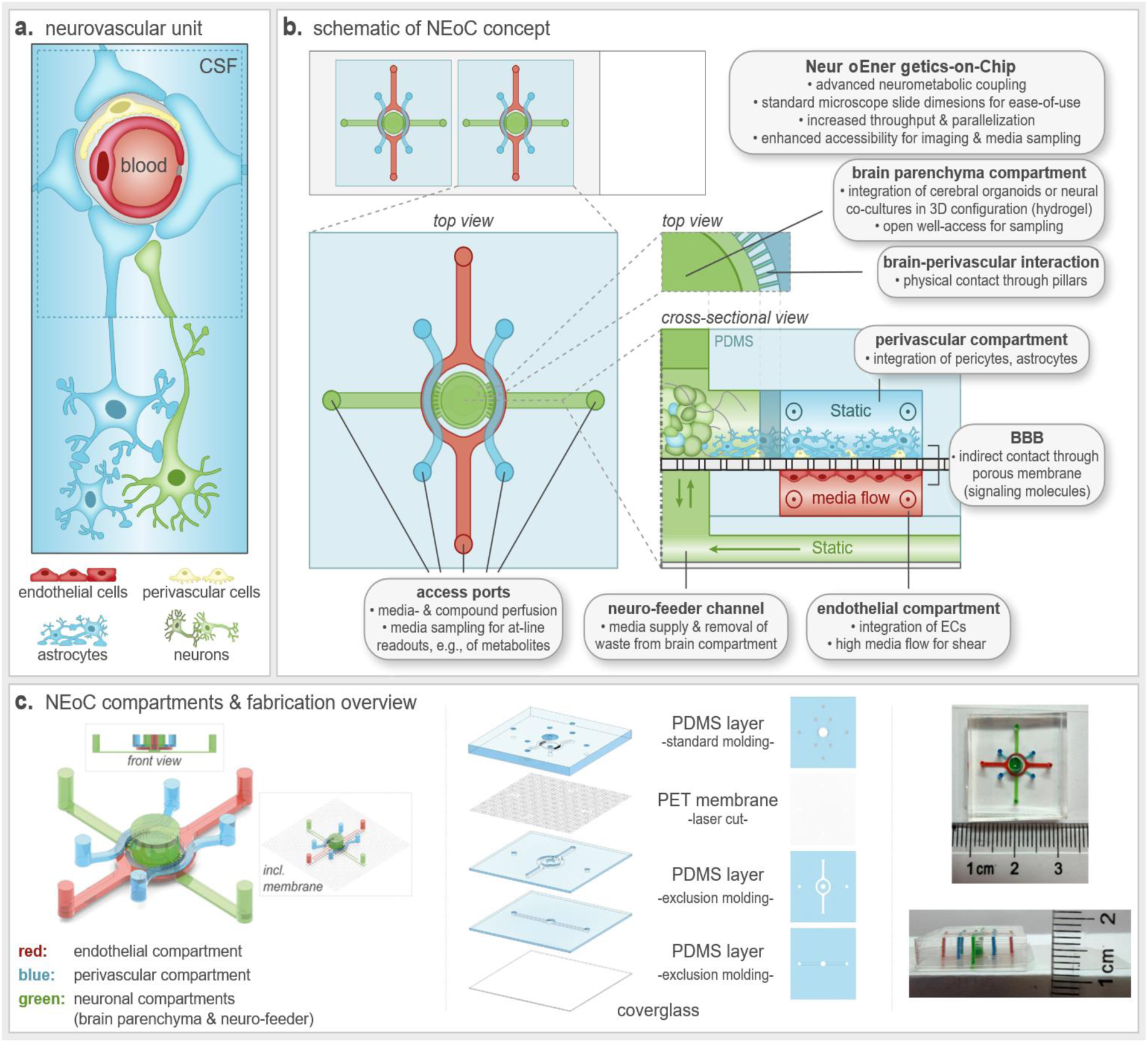

## Introduction

The human brain is a highly energy-demanding organ, accounting for approximately 20% of total body energy consumption [1]. This demand is supported by a specialized neurovascular network, the neurovascular unit (NVU), a highly organized multicellular system composed of endothelial cells, pericytes, astrocytes, microglia, neurons, and extracellular matrix components [2]. The NVU plays a central role in maintaining brain homeostasis by tightly regulating the exchange of nutrients, metabolites, and gases, which is essential for proper central nervous system (CNS) function [3–6]. Among these exchanged nutrients, glucose represents the brain’s principal energy source, and the delivery of its energy from the vasculature to neurons depends on coordinated transport and metabolism across each cellular component of the NVU.

Given the brain’s limited energy storage capacity, a continuous supply of glucose from the circulation is essential, and meeting neuronal energy demand requires coordinated handling of this glucose across multiple cell types of the NVU. Glucose transport is primarily mediated by facilitative transporters of the solute carrier family 2 (SLC2A): glucose transporter 1 (GLUT1) mediates uptake into endothelial cells and astrocytes, while glucose transporter 3 (GLUT3) mediates neuronal uptake [7, 8]. Astrocytes occupy a central position in this network, functionally linking the vascular and neuronal components, for instance, in a major neuron support mechanism known as the astrocyte-neuron lactate shuttle (ANLS): by taking up glutamate, astrocytes stimulate aerobic glycolysis, leading to increased glucose consumption and lactate release to support neuronal energy demands [4, 7, 8]. Beyond glucose, neurons and glial cells can also draw on alternative substrates, such as monocarboxylic acids and fatty acids, reflecting the brain’s broader metabolic flexibility [9]. Disentangling how these cell type-specific contributions combine into a functional metabolic network requires experimental models in which the relevant cell populations can be co-cultured, perturbed, and monitored individually.

Disruptions in these tightly regulated processes are associated with a wide range of neurological and neurodegenerative disorders, and altered brain metabolism has been implicated in conditions such as Alzheimer’s disease, traumatic brain injury, and ischemic stroke [9–11]. In particular, age-related declines in GLUT1, GLUT3, and GLUT4 expression impair glucose uptake, enhance the accumulation of reactive oxygen species (ROS), and contribute to progressive neuronal dysfunction [12]. Together, these observations establish metabolic dysregulation as both a hallmark and an early driver of CNS pathology, and a deeper, cell-resolved understanding of NVU metabolic coupling may inform future approaches to early diagnosis and intervention.

Capturing these processes experimentally, however, remains challenging as current models fail to fully recapitulate human brain energy metabolism. Traditional two-dimensional in vitro cultures lack the structural and functional complexity of the native tissue, while the translatability of animal models is often limited by interspecies differences [13]. To address these limitations, the use of human-based microphysiological systems (MPS) has expanded significantly over the past two decades [14]. Among these, microfluidic organ-on-chip (OoC) platforms have emerged as powerful tools for modeling human biology enabling precise control of microenvironmental conditions, including dynamic perfusion, regulated nutrient delivery, and real-time monitoring of cellular behavior [15, 16]. OoC platforms further support controlled cell-cell interactions, spatial organization, and tissue architecture through the use of microchannels, membranes, and compartmentalized structures [17], and their compatibility with human induced pluripotent stem cells (iPSCs) enhances their physiological relevance and enables patient-specific disease modeling [18, 19].

OoC NVU and blood-brain barrier (BBB) models have been shown to be promising for investigating cell interactions in the central nervous system as well as disease modeling and drug delivery, as they enable controlled recapitulation of neurovascular interfaces [20–23], and iPSC-derived NVU and BBB models represent a solid platform for personalized modeling in health and disease [24–26]. However, these models typically focus on the vascular and perivascular interface alone, without an integrated brain parenchymal compartment, limiting their ability to capture the full metabolic exchange between the vasculature and the neural tissue it supplies. The combination of OoC and stem cell technologies has great potential to assess neurometabolic coupling by evaluating glucose transport, energy substrate utilization, and BBB function across both the vascular interface and the adjacent brain parenchyma, while enabling precise manipulation of individual cellular components to dissect their metabolic contributions.

Here, we present a novel OoC platform that recapitulates human neurometabolic coupling, hereafter referred to as the NeuroEnergetics-on-Chip (NEoC) system. By integrating isogenic cerebral organoids with vascular, perivascular, and astrocytic components within a dynamic, compartmentalized, and perfused microenvironment, this system provides a robust and physiologically relevant model of human brain metabolism. To our knowledge, the NEoC platform is the first to incorporate a dual-sided BBB interface, with its perivascular compartment directly connected to an open-well organoid chamber via microchannels. This configuration enables close spatial proximity between cellular compartments, facilitating paracrine signaling and cell–cell interactions, while preserving independent control over fluidic conditions and nutrient supply. Using this platform, we demonstrate that NVU/brain parenchyma co-culture with cerebral organoids modulates the phenotype of endothelial, astrocytic and pericyte populations, and that glucose is transferred across compartments in a co-culture-dependent manner, accompanied by distinct shifts in the secreted metabolome. Collectively, the NEoC platform enables investigation of cell-type-specific contributions to metabolic regulation, supports studies of disease-associated alterations, and provides a translational framework for the development of targeted therapeutic strategies.

## Results & discussion

### Design and Novelty of the NEoC Microfluidic Platform

Our microfluidic platform introduces a previously unrealized integration of a classical barrier-on-chip architecture with a spatially adjacent, three-dimensional tissue/organoid culture module. To our best knowledge, this is the first system to combine an on-chip barrier component through a direct lateral, small-distance fluidic interface that preserves both intercellular communication and imaging accessibilty (**Figure 1**).

**Figure 1.**
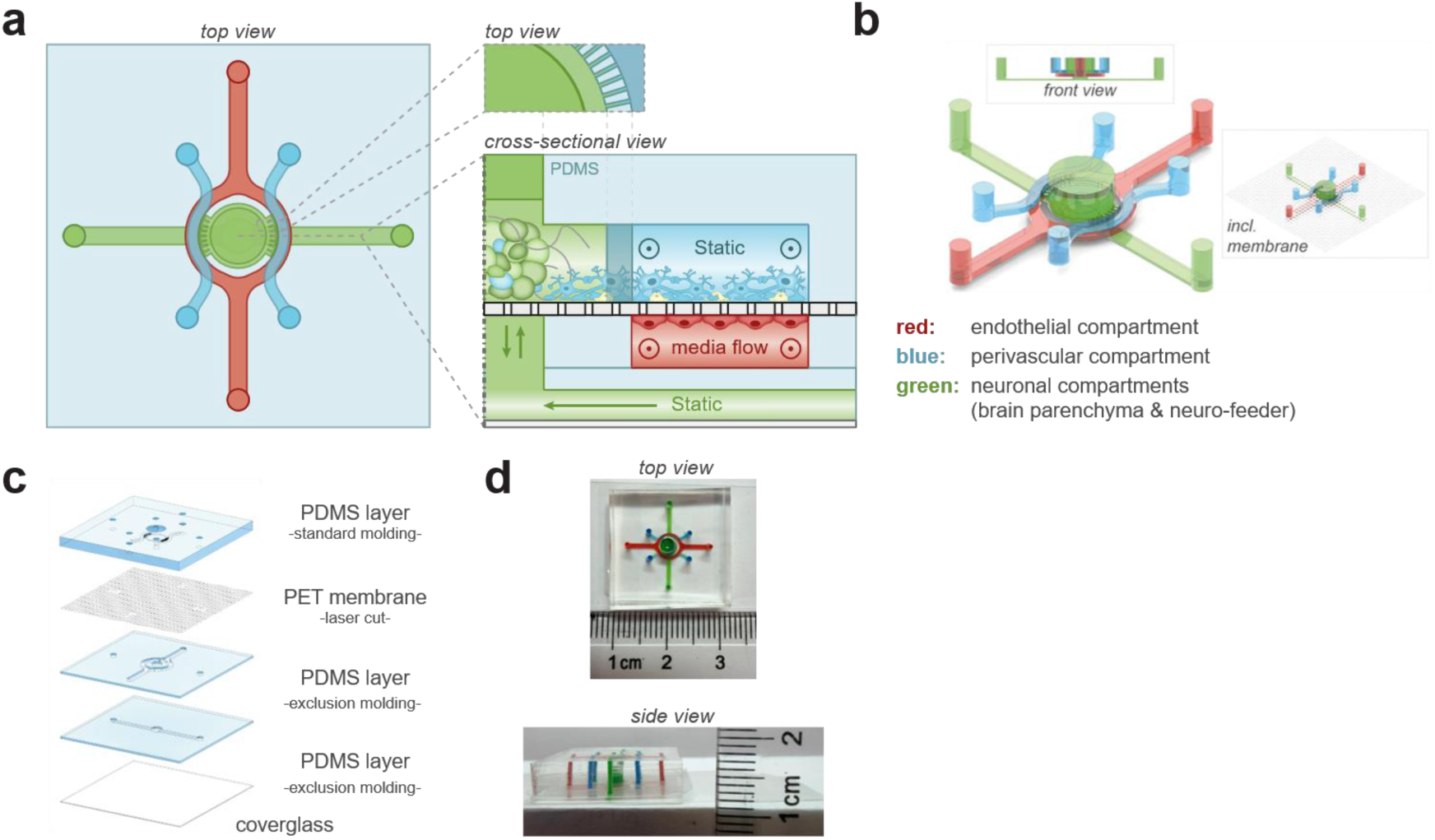
Design of the NEoC platform. **a.** Simplified schematic illustrating of the NEoC concept. **b.** Three-dimensional view of the three cellular compartments. **c.** Exploded-view schematic detailing the structural components of the microfluidic device. **d.** Representative photograph of an assembled NEoC device, with channels filled with colored liquids matching the compartment colors used in the schematics, to visualize the individual compartments..

The device comprises three sequential compartments – an endothelial channel, a perivascular channel and a brain parenchymal chamber – arranged to recapitulate the structural organization of the NVU while extending it into the adjacent brain tissue it serves. The endothelial compartment (**Figure 1**, in red) accomodates iPSC-derived endothelial cells seeded as a monolayer and aims to resemble the brain vasculature; medium is perfused through this channel to mimic blood circulation. Endothelial cells lining this compartment communicate with the perivascular compartment through an isoporous, semipermeable membrane that enable paracrine (and potentially endocrine) signaling across the two cell layers. The perivascular compartment (**Figure 1**, in blue) supports a co-culture of iPSC-derived pericytes and astrocytes seeded in a monolayer to the opposite side of this membrane, thereby completing the cellular constituents of the BBB. Critically, this compartment is not an endpoint: it connects directly to the parenchymal compartment via lateral microchannels, which permit not only the exchange of soluble factors but also physical cell-cell interactions, such as the outgrowth of cellular processes between the perivascular space and the brain parenchyma. Alongside the perivascular compartment, the top layer constitutes this parenchymal compartment (**Figure 1**, in green), comprising an open well that integrates 3D neural structures such as iPSC-derived cerebral organoids. Finally, to independently supply the parenchymal compartment with medium without disrupting the barrier-parenchyma interface, we incorporated a dedicated neurofeeder layer (**Figure 1**, green) that maintains contact with the parenchymal compartment via the membrane.

Each compartment maintains its own dedicated fluidic circuit, enabling distinct media formulations and flow regimens tailored to the physiological demands of each tissue type: continuous shear stress is applied exclusively to the endothelial channel; the perivascular chamber remains under static conditions; and the more demanding 3D parenchymal compartment is shielded from shear entirely, instead relying on diffusive exchange from the adjacent perfused neurofeeder channel. The entire device is mounted on a glass coverslip, providing full optical access for live microscopy across all three compartments.

Unlike the few existing barrier-organoid hybrid models, in which vertical stacking obscures imaging and complicates barrier-organoid interfacing, our lateral connection strategy preserves unobstructed optical access while maintaining stable, direct tissue-to-tissue integration between the BBB and the 3D parenchymal compartment. While demonstrated here for neuroenergetic studies, this architecture is inherently versatile and readily adaptable to any barrier tissue coupled to a 3D culture, ranging from extracellular-matrix-embedded single cells to organoids, spheroids, solid tumors, or ex vivo biopsies. This provides a broadly applicable tool for investigating barrier-3D tissue interactions, such as leukocyte diapedesis into solid tumor microenvironments, or the impact of an adjacent thrombus on barrier integrity and tissue viability in a stroke-like setting. The same approach could also be extended to other barrier types, such as the gut epithelium or placental trophoblast, to study nutrient and signaling exchange with their adjacent tissues.

### Microchannels enable diffusive exchange while limiting shear stress

To assess whether perfusion of the perivascular compartment could expose the adjacent parenchymal compartment to disruptive shear forces, we performed computational fluid dynamics simulations of the NEoC platform (**Figure 2**). While the perivascular compartment is operated under static conditions in this study, we modeled a perfused scenario at 1.3 µL/min, a flow rate representative of physiologically relevant perfusion in microfluidic NVU models. The resulting velocity field shows that flow remains confined to the perfused perivascular channel, with velocities dropping to near zero across the lateral microchannels and the parenchymal compartment. This indicates that, at this flow rate, the parenchymal compartment (and any 3D tissue cultured within it) remains shielded from shear stress, consistent with its intended role as a low-shear, diffusion-dominated microenvironment.

**Figure 2.**
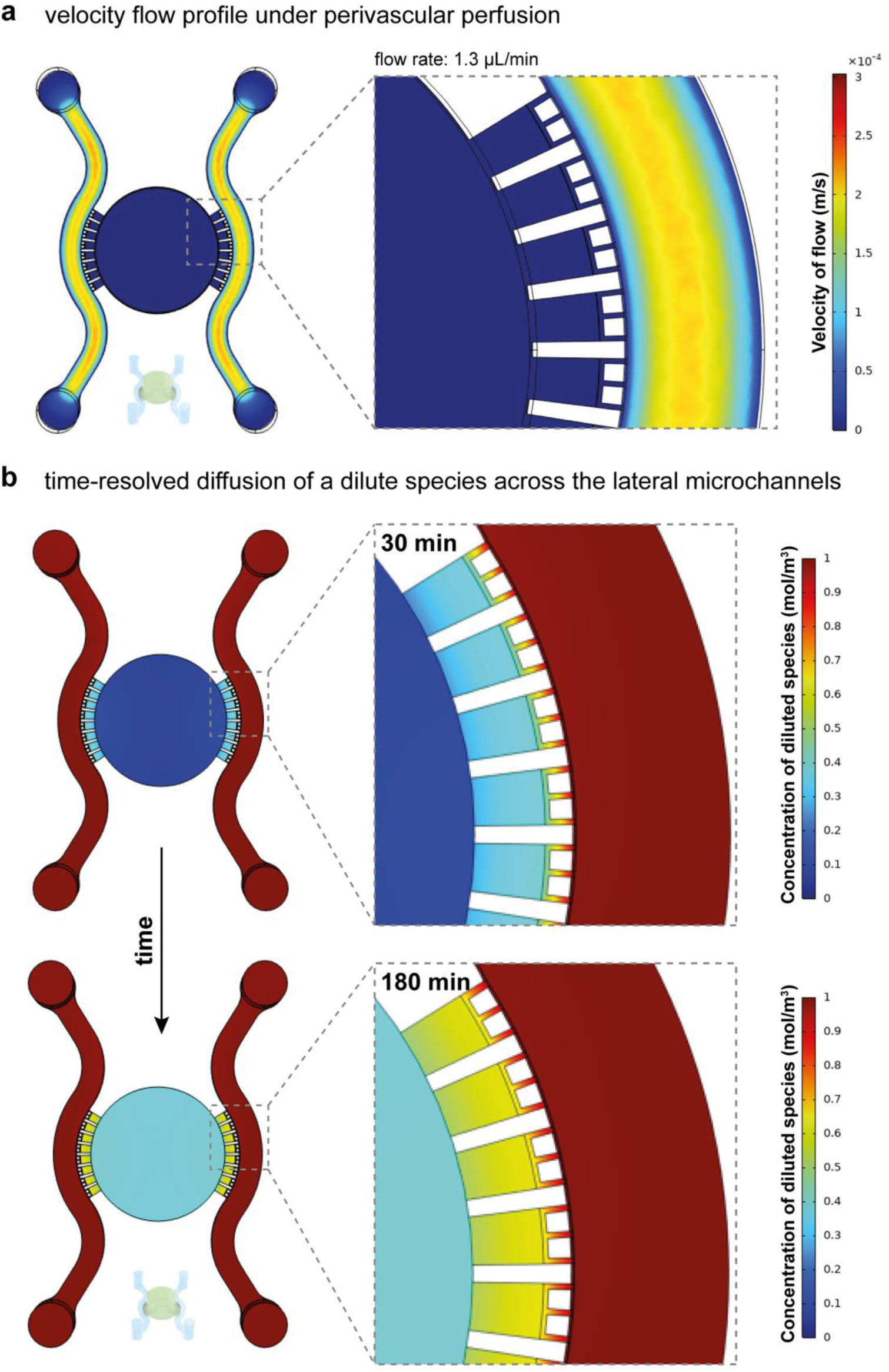
Computational modeling of flow and diffusion within the NEoC platform. **a.** Simulated velocity flow profile under perivascular perfusion at 1.3 µL/min. Flow is confined to the perfused perivascular channel, while the lateral microchannels and the central parenchymal compartment show negligible flow velocity, indicating that the parenchymal compartment is shielded from convective flow and shear stress. **b.** Time-resolved simulation of the diffusion of a dilute species (1 mol/m³ initial concentration in the perivascular channel) across the lateral microchannels and into the parenchymal compartment. At 30 min, the concentration front has progressed partway through the microchannels; by 180 min, the diluted species has diffused through the microchannels and begins to accumulate within the parenchymal compartment, demonstrating diffusive exchange between compartments in the absence of convective flow.

While the lateral microchannels restrict convective flow, they remain open for molecular exchange between compartments. To evaluate this, we simulated the diffusion of a dilute species from the perivascular channel into the parenchymal compartment over time (**Figure 2b**). Within 30 minutes, the diffusion front had progressed partway through the lateral microchannels, and by 180 minutes the diluted species had traversed the full length of the microchannels and begun to accumulate within the parenchymal compartment. These results indicate that, despite the absence of convective transport, the lateral microchannel geometry permits diffusive communication between the perivascular and parenchymal compartments on a timescale of hours – supporting paracrine signaling and nutrient exchange without compromising the static, low-shear environment required for 3D tissue culture.

### Profilometry confirms high-fidelity replication of microfluidic features

The PDMS layers of the NEoC were fabricated by replica molding using HTL-20 resin master molds (see Supplementary Methods for details). To assess the fidelity of feature transfer from the resin mold to the PDMS layers, three-dimensional optical profilometry was performed on both structures. The endothelial channel layer showed high replication accuracy; both the master mold and the corresponding PDMS layer exhibited comparable maximum heights of approximately 200 µm (**Figure 3a-b**). Further measurements confirmed that the channel height remained consistent along its length, as measured at three representative cross-sectional positions (**Figure 3c-e**). Maintaining this channel height is important for ensuring reproducible shear stress conditions across the endothelial monolayer, as even small deviations in channel geometry can alter local flow velocity and the resulting wall shear stress experienced by the cells.

**Figure 3.**
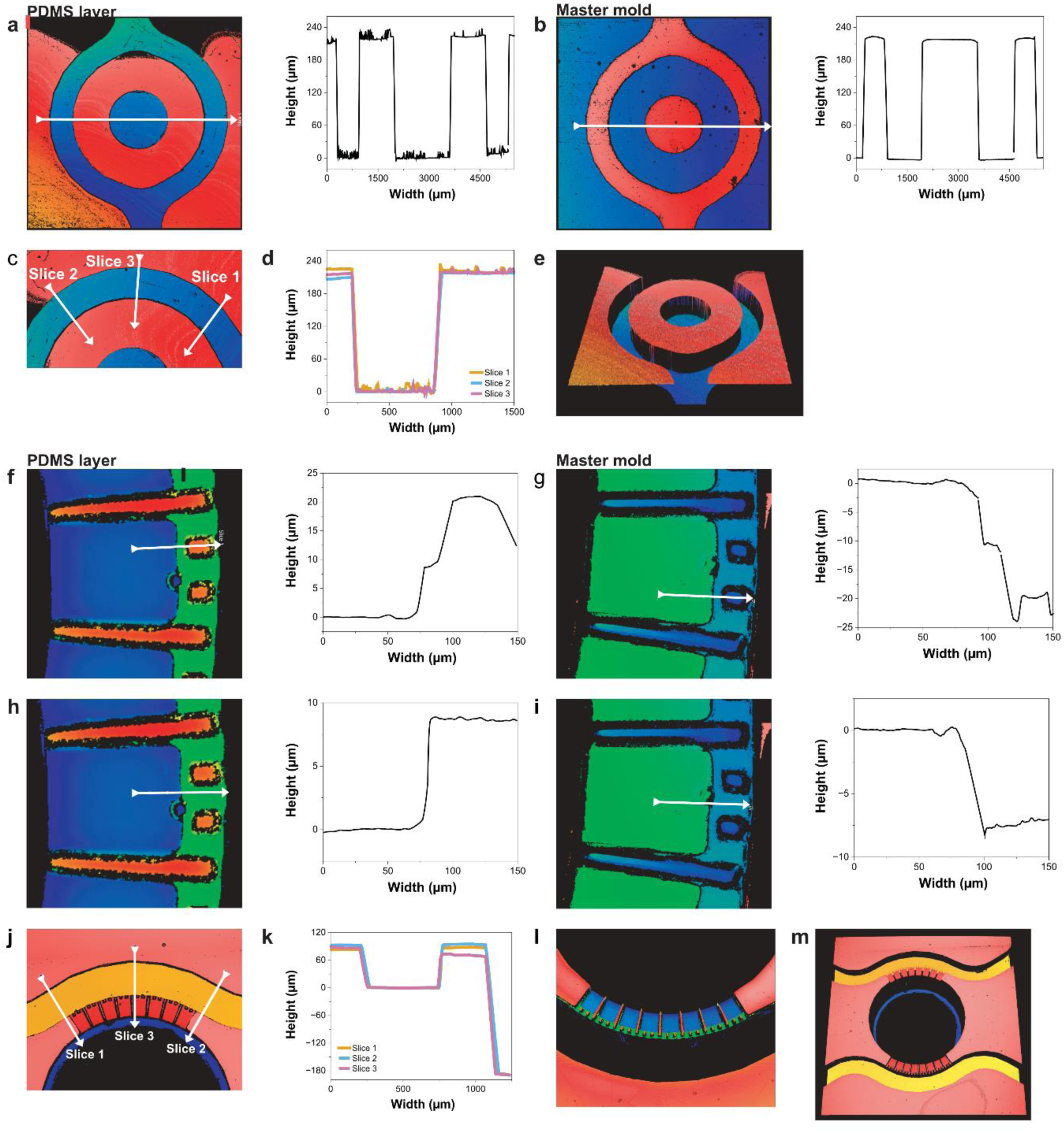
Three-dimensional surface profiles of the endothelial compartment and the top layer comprising the perivascular and the parenchymal compartments acquired by 3D optical profilometry. **a**. Surface profile of the endothelial channel PDMS layer after fabrication, with the corresponding graph showing the channel dimensions. **b**. Surface profile of the master mold used for fabrication of the PDMS layer. **c-d**. Channel dimensions measured at three representative cross-sectional positions along the endothelial channel PDMS layer, confirming dimensional consistency along its length. **e**. 3D surface profile of the PDMS layer. **f-i.** Surface profile of microchannels in the top layer: PDMS vs master mold. **j-k.** Comparison of dimensions across three cross-sectional slices of the perivascular channel: slice 3 spans a microchannel, confirming the expected height difference between the microchannels and the perivascular channel. **l-m.** 3D visualization of the top layer. The open well structure of the parenchymal compartment is shown in black; the microchannels are shown in green; the punching margins are shown in blue; the micropillars between the microchannels are shown in red. Color map represents elevation (warm colors: highest; cool colors: lowest).

A similar characterization was performed for the top PDMS layer, which contains the perivascular channel and the microchannel network connecting to the parenchymal compartment. This layer incorporates more intricate microstructures that are critical for maintaining spatial compartmentalization between the perivascular and parenchymal cell populations. Profilometry measurements indicated that the intended height differences between the main channels, connecting microchannels, and well-like structures were preserved. Comparison with the corresponding master molds confirmed a high degree of structural fidelity in the replicated PDMS features (**Figure 3f-l**). Quantitative analysis further demonstrated uniformity in the dimensions of micropillars and microchannels, with heights of approximately 12 µm for microchannels and 20 µm for micropillars (**Figure 3j-h, S3**). Together, these results indicate that the replica molding process reliably reproduces the layered height profile required to maintain distinct fluidic regimes across compartments and support the reproducibility of the NEoC platform across independent fabrication batches.

### Integration and characterization of hiPSC-derived NVU components in the NEoC platform

Despite their informative value, existing NVU models remain simplified, addressing only the BBB component, lacking parenchymal tissue, and thus limiting their ability to reproduce the complexity of human physiology and microenvironment [24, 27–29]. Different cell sources have been explored to model the NVU, each with distinct trade-offs: human primary brain cells, which offer a close resemblance to physiology but are limited in availability and exhibit batch-to-batch variability; immortalized cell lines are readily available but, being derived from or transformed by oncogenic processes, may exhibit, amongst others, aberrant metabolic profiles that poorly reflect the energetic demands of healthy brain tissue. Human iPSCs overcome both limitations, offering a scalable, renewable source that retains key physiological characteristics of the target cell types [19, 22]. Crucially, deriving all NVU cell types from a single isogenic iPSC line eliminates the interfering effect of donor-to-donor genetic variability, ensuring that observed differences in cell behavior reflect the experimental conditions rather than underlying genetic heterogeneity. This is an important consideration, especially when studying metabolic crosstalk across multiple, interacting cell populations.

To establish the NEoC model, hiPSCs were differentiated into CD31⁺ endothelial-like cells, CD31⁻ perivascular-like cells, and astrocytes (**Supplementary Fig. 4a–c**). Cell stocks of each cell type were cryopreserved and thawed as needed for integration into the microfluidic platform. Human iPSC-derived cerebral organoids were generated 20 days prior to chip loading, while CD31^+^-, CD31^-^-, and astrocyte cell populations were thawed and expanded in parallel, 10 days prior to chip loading. All cell types were subsequently harvested and introduced into the microfluidic device in a sequential seeding protocol: on day 0, cerebral organoids were encapsulated in Matrigel hydrogel and pipetted into the open well, followed by endothelial cells seeded into the endothelial compartment and allowed to adhere to the membrane with the chip inverted. To prevent Matrigel from blocking the perivascular channel, the channel was flushed prior to full polymerization. On day 1, a mixture of pericyte-like cells and astrocytes was introduced to the perivascular compartment. A schematic overview of the cell integration and on-chip culture workflow is presented in **Figure 4a**.

**Figure 4.**
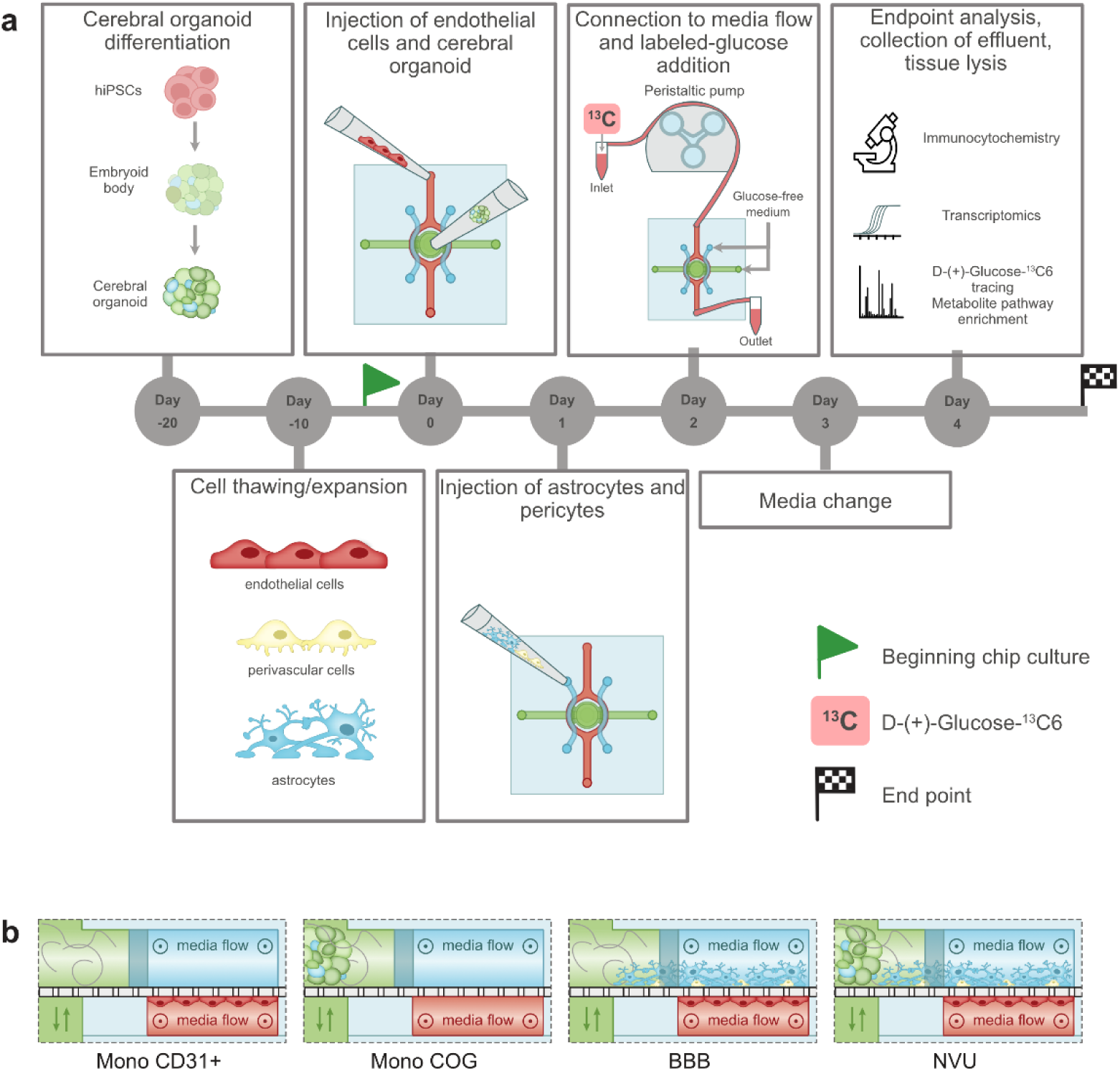
Integration and biological characterization of isogenic hiPSC-derived NVU components in the NEoC. **a.** Schematic overview of NEoC experiment timeline. **b.** Experimental configurations tested.

To characterize the behavior of each cell population within the NEoC platform, four experimental configurations were evaluated (**Figure 4b)**:

i. CD31⁺ cells in monoculture (*Mono CD31^+^*),
ii. cerebral organoids in monoculture (*Mono COG*),
iii. co-culture of CD31⁺ cells, astrocytes, and CD31⁻ cells (*BBB*), and
iv. full co-culture of all four populations, including CD31⁺ cells, astrocytes, CD31⁻ cells, and cerebral organoids (*NVU*).

For each condition, five chips were run in parallel per experiment, alongside one acellular device as a negative control.

Endothelial cells play a central role in maintaining brain homeostasis by regulating the transport of molecules across the blood-brain barrier through tightly regulated cell-cell junctions. These junctions are mediated by adhesion proteins such as VE-cadherin and PECAM-1 (CD31). Within the NEoC platform, CD31⁺ cells exhibited increased PECAM-1 expression in the *NVU* condition, whereas expression was reduced in both the *monoculture* and *BBB* conditions compared to 2D controls (**Figure 5a**). In contrast, VE-cadherin expression was elevated in both *BBB* and *NVU* conditions relative to monoculture and 2D controls, suggesting that interactions with additional cell types enhance endothelial phenotype maturation (**Figure 5b**), as previously reported [30, 31]. The variability observed in the *NVU* condition, reflected by the large standard error, likely captures the inherent biological complexity of a fully assembled multicellular system, in which organoid-to-organoid differences in maturation state and size may introduce condition-specific variance not present in simpler configurations.

**Figure 5.**
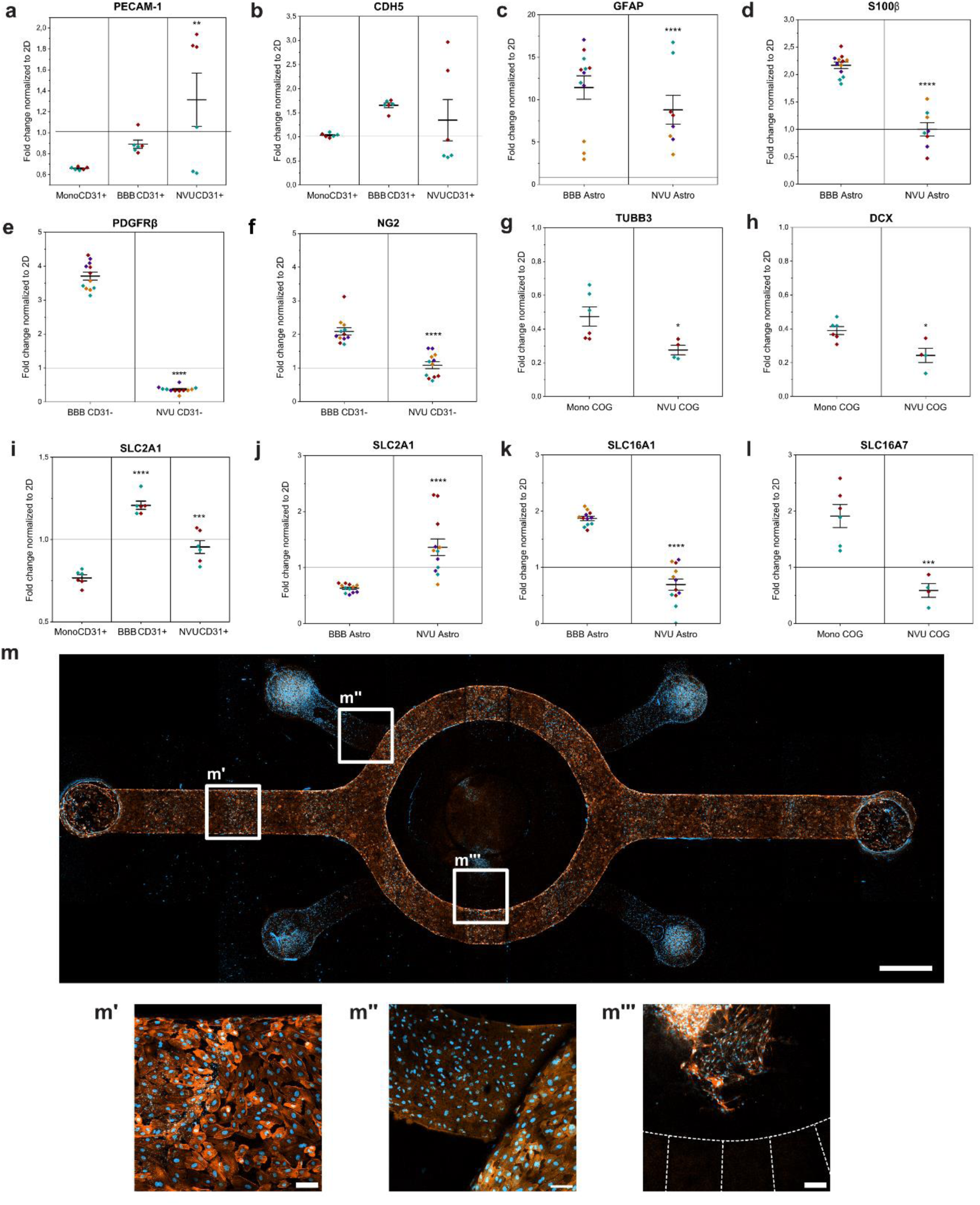
Gene expression analysis and special organization of hiPSC-derived NVU components in the NEoC. a-l. mRNA expression analysis of hiPSC-derived cell populations across the different NEoC culture configurations on day 4 of chip culture. Data represented as fold change normalized to the corresponding 2D culture controls. Whiskers indicate ± SE around the mean (bar). Fold change > 1: upregulation of the gene, fold change <1: downregulation, and fold change = 1: unchanged expression. The color of each data point indicates the biological replicate (chip), analyzed in duplicate or triplicate. P-values were derived using linear mixed models (LMM) *p < 0.05, **p < 0.01, ***p < 0.001, ****p < 0.0001. **a-b**. PECAM-1 and VE-Cadherin (CDH5) expression from CD31+ cells, p-value comparison versus Mono CD31^+^, n=2. **c-f.** GFAP, S100β, PGFRβ, and NG2 expression in co-cultures of astrocytes and CD31^-^ pericyte-like cells; p-value comparisons versus BBB condition, n=4. **g-h.** TUBB3 and DCX expression in cerebral organoids, p-value comparison versus Mono COG, n=2. **i-j.** SLC2A1 (encoding GLUT1) expression comparison in CD31^+^ cells (**i**) or in astrocytes (**j**), p-value comparison versus Mono CD31^+^ or BBB astrocytes, respectively. **k.** SLC16A1 (encoding MCT1) expression in astrocytes, p-value comparison versus BBB astrocytes, n=4. **l.**SLCA16A7 (encoding MCT2) expression in cerebral organoid, p-value comparison versus Mono COG, n=2. **m.** Representative immunofluorescence image of the NVU condition, showing the spatial distribution of all cell populations across the microfluidic device. Nonspecific DAPI signal was removed for clarity (Figure S4a). Orange (phalloidin), blue (DAPI). Scale bar equals 1000 µm. **m’.** Representative image of an endothelial channel. **m’’.** The perivascular channel (top left) and endothelial channel (bottom right) showing the compartmental interface. **m’’’.** Cerebral organoid in the parenchymal compartment extending projections toward the microchannels. Orange (phalloidin), blue (DAPI). Scale bars equal 100 µm.

Astrocytes are essential components of the NVU and brain tissue in general; they contribute to CNS regulatory function, particularly by sensing and controlling glucose transport across the BBB and by producing alternative energy substrates, such as lactate, as described in the ANLS framework [7]. Pericytes, located perivascularly around brain endothelial cells of the BBB, play a vital role in regulating vessel maturation, stabilization, and endothelial barrier function and proliferation [32]. Together, these two cell populations form the perivascular niche of the BBB, and their response to the broader multicellular environment of the NEoC provides a sensitive readout of how the presence of brain parenchymal tissue shapes the perivascular phenotype – and vice versa.

We co-cultured iPSC-derived astrocytes and pericyte-like cells within the perivascular compartments of the NEoC platform. Astrocytes cultured in the *BBB* condition showed more than a 10-fold increase in GFAP expression compared to 2D controls (**Figure 5c**); however, in the *NVU* condition, GFAP expression decreased compared to the *BBB* configuration. Rather than reflecting reduced astrocytic engagement, this pattern is consistent with a shift toward a more homeostatic, less reactive astrocyte state in the presence of neural tissue, as reported in previous studies with iPSC-derived astrocytes co-cultured with neurons [33]. In contrast, S100β expression in astrocytes was reduced in the *NVU* condition, reaching levels comparable to those observed in 2D culture (**Figure 5d**). Since elevated S100β expression has been associated with astrocyte activation and astrocytic inflammatory signaling [34–36], its reduction in the *NVU* condition suggests that the presence of the cerebral organoid attenuates astrocytic stress responses, a finding consistent with the GFAP trajectory and indicative of a more physiologically balanced astrocyte phenotype when neural tissue is present.

CD31⁻ pericyte-like cells similarly displayed differential marker expression across different culture conditions: PDGFRβ expression was decreased in the *NVU* condition but increased more than 3-fold in the *BBB* condition relative to 2D control levels. A similar trend was observed for NG2 expression (**Figure 5e-f**). Since PDGFRβ and NG2 are established markers of pericyte identity and vascular stabilization function, these findings suggest that the presence of the cerebral organoid modulates pericyte marker expression in a manner that may reflect altered paracrine signaling demands within the perivascular compartment. Collectively, the gene expression patterns across both astrocytes and pericyte-like cells indicate that the presence of cerebral organoids shapes perivascular cell phenotype, highlighting the importance of multicellular crosstalk between the brain parenchyma and the perivascular niche for NVU homeostasis.

Neural activity accounts for the major energy expenditure in the central nervous system. Neurons communicate their energy needs to their surroundings, such as glia and the neurovasculature, which, in turn, provide the energy substrate. To mimic the neuronal component in the NEoC, we integrated iPSC-derived cerebral organoids at day 20 of differentiation into the parenchymal compartment of the microfluidic device. Expression of the neuronal markers βIII-tubulin (TUBB3) and doublecortin (DCX) was decreased in about 0.2 to 0.4-folds in both the monoculture and *NVU* conditions (**Figure 5g-h**). This, for once, likely reflects the well-documented inter-organoid variability inherent to unguided differentiation protocols, which can substantially influence gene expression readouts, particularly at early time points. Moreover, the transition into the microfluidic environment, with its distinct nutrient gradients compared to standard suspension culture, may impose additional constraints on the cerebral organoids. Optimization of organoid integration and culture conditions represents an important next step for future iterations of the NEoC system. Notably, cerebral organoids remained structurally intact and extended cellular processes toward the microchannels (**Figure 5m’’’**), confirming that the parenchymal compartment supports viable 3D neural tissue and demonstrating active engagement with the surrounding microenvironment.

The cellular components of the NVU are metabolically coupled through coordinated glucose and monocarboxylate transport processes, that supply neurons with energy substrates. Glucose is transported from the bloodstream into endothelial cells via GLUT1, then taken up by astrocytes, and metabolized to lactate via aerobic glycolysis. Lactate is then exported by astrocytes via Monocarboxylate Transporter 1 (MCT1) and taken up by neurons through MCT transporters, representing one proposed mechanism of metabolic coupling between astrocytes and neurons [7].

To assess whether metabolic coupling is established in the NEoC model, we examined the expression of key transporters involved in this process. In CD31⁺ endothelial cells, GLUT1 expression was higher in both *BBB* and *NVU* conditions compared with monoculture, suggesting that the co-culture environment promotes glucose uptake capacity in endothelial cells; however, levels in the NVU condition remained lower than those observed in 2D controls (**Figure 5i**). This may reflect the distinct culture medium compositions and substrate availability in the chip environment compared to 2D conditions, rather than a fundamental impairment of endothelial metabolic function. In astrocytes, GLUT1 expression was markedly increased in the *NVU* condition compared with both *BBB* and 2D cultures, suggesting enhanced metabolic activity in the multicellular environment (**Figure 5j**). This upregulation is consistent with astrocytes increasing their glucose uptake capacity in response to signals from the adjacent neural tissue, supporting the notion that the NEoC captures physiologically relevant metabolic crosstalk between the parenchymal and perivascular compartments. Given that neurons predominantly express the glucose transporter GLUT3, we also assessed its expression in organoids to evaluate their reliance on glucose metabolism. GLUT3 expression showed a trend toward elevation under NVU conditions, which would be consistent with increased neuronal metabolic demand in the presence of the full NVU complement; however, the high variability in the data precludes confident conclusions (F**igure S4**).

Alongside these glucose transport findings, we examined the expression of the monocarboxylate transporters MCT1 in astrocytes and MCT2 in cerebral organoids, as readouts of lactate export and import capacity, respectively. MCT1 expression in astrocytes was reduced under *NVU* conditions (**Figure 5k**), and MCT2 expression in cerebral organoids was similarly decreased in the *NVU* condition (**Figure 5l**). This downregulation of both transporters in the most complex culture condition might reflect the inherent transcriptional variability of the system, complicated by the inter-organoid variability. Notably, the concurrent upregulation of astrocytic GLUT1 in the *NVU* condition confirms that the perivascular compartment is metabolically responsive to the multicellular environment. Investigating lactate-based metabolic coupling more conclusively will be an important focus as organoid maturation and on-chip culture conditions are further optimized.

To confirm the successful integration and spatial organization of each cell population within their intended compartments, we performed immunofluorescence imaging of the assembled NEoC platform. We achieved a homogeneous distribution of CD31⁺ cells within the chip, where staining revealed a continuous, confluent cell layer indicating barrier formation (**Figure 5m’**). The interface between the perivascular and endothelial channels is visible in **Figure 5m’’**, confirming compartmental organization and spatial separation of the two cell populations during co-culture. Cerebral organoids within the parenchymal compartment extended projections toward the microchannels (**Figure 5m’’’’**), suggesting active interactions between parenchymal and perivascular compartments. Further immunofluorescence staining for VE-cadherin, PDGFRβ, and S100β confirmed the presence of the respective cell populations within their intended compartments, with cell-type identity supported by compartmental location (**Figure S5**). Together, these observations confirm the successful spatial integration of all NVU cell populations within the NEoC and provide the structural basis for the intercellular interactions further characterized below.

A central strength of the NEoC platform lies in its use of hiPSCs to generate all NVU cell types from a single isogenic source, enabling physiologically relevant, patient-specific modelling of the neurovascular-to-brain parenchyma interface. To capture the structural and functional complexity of the human brain parenchyma, we incorporated self-organized cerebral organoids within the parenchymal compartment as the primary neuronal model in this study. However, the open-well architecture of the parenchymal compartment is not limited to organoids, but it is designed to accommodate a range of 3D neuronal constructs. To demonstrate this, we evaluated the integration of alternative neural sources, including single neurons and neurospheres derived from neuroectodermal stem cells, both of which where successfully maintained within the parenchymal compartment (**Figure S6**). These pilot experiments confirm that the NEoC platform is compatible with diverse 3D neural models, broadening its applicability beyond the cerebral organoid system demonstrated here. Combined with the use of patient-derived iPSCs, this flexibility positions the NEoC as a versatile tool for investigating disease-specific neurovascular and neurometabolic phenotypes across a range of neural model systems and pathological contexts.

### Glucose transfer and metabolic profiling demonstrate compartment-dependent metabolic coupling

Understanding how glucose moves from the vasculature through the BBB and into the brain parenchyma is central to studying neurometabolic coupling, and disruptions in this process are implicated in a range of neurological disorders. To investigate this process within the NEoC, we evaluated glucose diffusion across the compartments of the microfluidic device under physiologically relevant, normoglycemic conditions. ^13^C-labeled glucose was introduced into the endothelial channel at 5.3 mM, while the perivascular and neurofeeder channels, as well as the parenchymal compartment, were maintained in glucose-free medium to enable unambiguous tracking of glucose transport from the vascular to the parenchymal compartment. After 48 hours, media samples were collected from all compartments, and the proportion of ^13^C-labeled glucose relative to total glucose was quantified with targeted mass spectrometry (MS).

Approximately 10% of the glucose detected in the perivascular compartment was ^13^C-labeled across all conditions (*mono-COG*, *BBB*, and *NVU*), confirming that labeled glucose successfully crossed from the endothelial channel into the adjacent perivascular compartment regardless of culture configuration. In contrast, within the parenchymal compartment, only ∼4% of glucose was labeled in the *mono-COG* condition, whereas higher proportions (∼7–9%) were observed in the *BBB* and *NVU* conditions (**Figure 6a**). This difference suggests that the presence of vascular and perivascular cell layer facilitates glucose transfer toward the parenchymal compartment – consistent with the known role of the BBB actively governing substrate trafficking between blood and brain parenchyma. While the underlying mechanisms and precise directionality of transport cannot be definitively established from these data alone, to co-culture-dependent increase in parenchymal glucose supports the physiological relevance of the NEoC.

**Figure 6.**
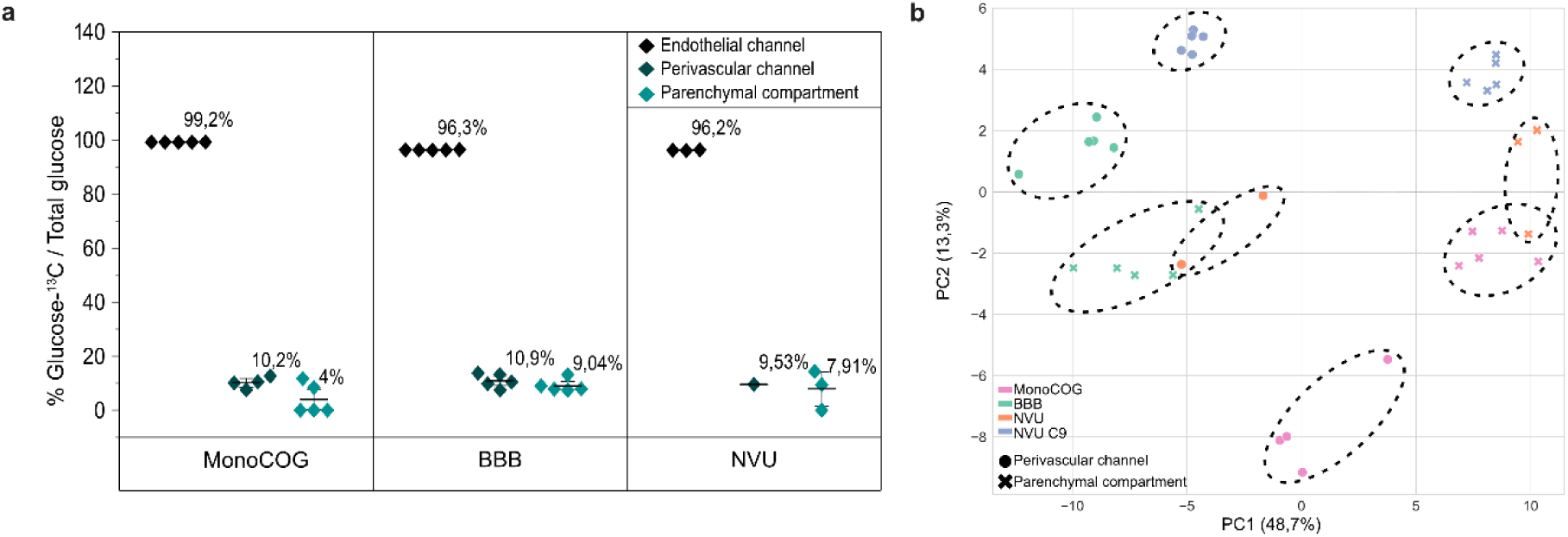
Glucose transfer and metabolic profiling across the NEoC compartments. **a.** Proportion of ^13^C-labeled glucose relative to total glucose quantification in each compartment of the NEoC, quantified by targeted mass spectrometry following 48 h of ^13^C-glucose perfusion through the endothelial channel. Each data point represents one biological replicate (chip), n=5. **b.** Unsupervised principal component analysis (PCA) of the compartment secretomes. The feature matrix (samples × features) was transposed and decomposed using singular value decomposition as implemented in scikit-learn. The percentage of variance explained by each principal component is reported on the respective axis.

Building on these findings on the level of glucose, we next sought to characterize the broader metabolic landscape of the NEoC. Alterations in metabolite levels may reflect changes in brain metabolism and could potentially contribute to early diagnostic approaches and to a better understanding of neuropathological processes [37]. To this end, we performed untargeted MS-based metabolomic profiling of the secretome collected from the perivascular and parenchymal compartments under *Mono-COG*, *BBB*, and *NVU* conditions, as well as an additional *NVU* condition generated from an independent apparently healthy iPSC line (NVU C9), to begin assessing inter-line reproducibility.

Principal component analysis (PCA) of the secretome revealed a clear separation between perivascular and parenchymal compartment profiles along PC1, which accounted for 48.7% of the total variance, confirming that the two compartments maintain metabolically distinct entities (**Figure 6b**). Within the parenchymal compartment, the *BBB* condition (in which no cells are present in the parenchymal compartment and detected metabolites primarily reflect diffusion from the perivascular space) clustered closely with the perivascular profiles, providing an important baseline for interpreting condition-specific parenchymal signals.

In contrast, the *Mono-COG* condition diverged markedly from this baseline, reflecting the organoid’s own metabolic contribution to the parenchymal secretome in the absence of perivascular influence. Notably, the *NVU* parenchymal profile occupied an intermediate position between the *BBB* and *Mono-COG* conditions, consistent with the organoid’s metabolic activity being modulated by signals diffusing from the perivascular compartment providing the first evidence of bidirectional metabolic communication between the vascular and parenchymal components of the NEoC. Notably, while the NVU C9 line clustered distinctly from the C14-derived *NVU* condition, both remained clearly separated from the parenchymal profiles, suggesting that platform-level metabolic compartmentalization is preserved across iPSC lines despite donor-specific metabolic variation.

## Conclusion

The NEoC platform presented here represents a significant step toward physiologically relevant, human-based modeling of neurovascular metabolic coupling. By integrating isogenic hiPSC-derived endothelial-like, pericyte-like, and astrocytic cells with cerebral organoids in a compartmentalized, perfused microfluidic device, we established, to our best knowledge, the first platform to combine a dual-sided BBB interface with a directly connected open-well organoid compartment, enabling simultaneous investigation of barrier function and parenchymal metabolic activity within a single, optically accessible device.

Biological characterization demonstrated that co-culture with cerebral organoids modulates perivascular cell phenotype in a manner consistent with a more homeostatic NVU state. Glucose tracing confirmed co-culture-dependent transfer of labeled glucose from the vascular to the parenchymal compartment under normoglycemic conditions, and untargeted metabolomic profiling revealed co-culture-driven changes of the metabolic signatures of the distinct cellular compartments.

As a proof-of-concept study, the findings presented here establish the NEoC as a functional platform for investigating neurovascular metabolic coupling, while naturally pointing toward areas for further development. Optimization of organoid maturation protocols and increased experimental throughput will strengthen the statistical resolution of individual readouts in future iterations. Looking ahead, longer culture periods and more mature organoids are expected to enable functional investigation of metabolic coupling mechanisms not yet fully established here, and the platform’s compatibility with patient-derived iPSC lines opens direct avenues for disease modeling and personalized therapeutic screening in neurological conditions characterized by metabolic dysregulation.

In summary, the NEoC bridges the gap between simplified in vitro systems and the complexity of human neurovascular biology, providing a versatile, human-specific tool for studying neurometabolic coupling in health and disease, with broad potential to accelerate the development of targeted therapeutic strategies for neurological disorders.

## 2. Materials and Methods

### 2.1 Microfluidic platform fabrication & characterization

The NEoC microfluidic platform was designed using Autodesk AutoCAD 2024. Key dimensions and compartment volumes are detailed in Supplementary Table 1.

#### 2.1.1 NEoC Chip fabrication

The NEoC microfluidic platform is a custom-designed, five-layer hybrid device consisting of three micropatterned polydimethylsiloxane (PDMS) layers bonded to a 24x60 mm rectangular coverglass (thickness 0.16-0.19 mm; #2980-246, Corning^®^) and separated by a semipermeable isoporous polyethylene terephthalate (PET) membrane (ipCELLCULTURE^TM^ Track Etched PET12; #2000M12/620M103, it4ip S.A). The PDMS layers were produced by replica molding using PDMS (mixed at a 10:1 (w/w) ratio of base polymer to curing agent; Sylgard 184 Silicone Elastomer Kit, Dow Corning) cast against 3D-printed, micropatterned master molds, which served as positive templates for feature transfer. A detailed fabrication process, including a schematic of the individual production and assembly steps (Figure S1), is provided in the Supplementary Information.

#### 2.1.2 Profilometry

Three-dimensional surface profiles of the micropatterned PDMS layers and master molds were acquired using a 3D optical profilometer (Nexview NX2, Zygo Corporation, USA). Measurements were performed using 2.75× and 10× objectives with 0.5×-2× zoom, selected based on the required lateral resolution (0.41-5.91 µm) and feature dimensions of the respective sample. Vertical scan lengths of 145-350 µm were applied depending on sample height. Surface topographies were assessed using Zygo Mx software, and quantitative data were subsequently exported and analyzed using custom Python scripts.

#### 2.1.3 Computational fluid dynamics modeling

Fluid flow and diffusive transport within the NEoC were modeled using COMSOL Multiphysics (COMSOL Version 6.2, Stockholm, Sweden). Incompressible, stationary free fluid flow was described by the Navier-Stokes equations, assuming the properties of water (dynamic viscosity μ = 1 × 10⁻³ Pa·s, density ρ = 1000 kg/m³) at a volumetric flow rate of 1.3 µL/min applied at the inlet of the perivascular channel. The geometry was reconstructed from the CAD design of the NEoC platform, capturing the perivascular channel, lateral microchannels, and parenchymal compartment. No-slip boundary conditions were applied at all channel walls, and a zero-pressure condition was imposed at the outlet.

Transport of a diluted species across the lateral microchannels and into the parenchymal compartment was modeled using the time-dependent convection-diffusion equation, with a diffusion coefficient of 1 × 10⁻⁹ m²/s and an initial concentration of 1 mol/m³ imposed at the perivascular channel inlet. Transport across the PET membrane was not included in the model. Simulations were run for 30 and 180 minutes to capture the temporal evolution of species diffusion into the parenchymal compartment.

### 2.1 NEoC Cell Integration

#### 2.2.1

hiPSC maintenance, differentiation, and expansion are detailed in Supplementary Information.

#### 2.2.2 On-chip Culture

The day before cell integration, chips were sterilized by exposure to O_2_-plasma (30 s, 50 W) andall channels were immediately filled with DPBS^−^ (Thermo Fisher Scientific, Cat. No. 37350) under sterile conditions. Chips were then centrifuged at 1000 × g for 10 minutes to facilitate degassing, and subsequently kept in DPBS^−^ to allow evacuation of residual air. Successful bubble removal was confirmed by bright-field microscopy.

The following protocol describes the full *NVU* condition, representing the most complex culture configuration. For simpler conditions (*Mono-CD31⁺*, *Mono-COG*, and *BBB*), chips were handled identically, with the exception that only the relevant compartments were seeded with cells. Matrigel coating was performed in all cases regardless of condition, and in the absence of a cerebral organoid, the parenchymal compartment was filled with Matrigel alone to maintain consistent hydrogel conditions across all configurations.

On day 0, the endothelial and perivascular channels were coated with Matrigel at a concentration of 0.084 mg/mL in DMEM-F12 and incubated for 1 hour at 37°C in a humidified incubator with 5% CO₂.

Following incubation, cerebral organoids were prepared for seeding by carefully removing residual Matrigel using a sterile needle. Then, 25 µL of Maturation Medium was added to the parenchymal compartment of the microfluidic device, and one organoid per chip was transferred, using a cut 1000 µL pipette tip, and allowed to settle to the bottom of the well. The medium was then carefully aspirated and replaced with 15 µL of Matrigel. The devices were incubated for 1 hour to allow Matrigel polymerization.

Subsequently, CD31^+^ cells were detached with 1X TrypLE (Cat. No. 12563011, Thermo Fisher), counted, and seeded at a density of 1.2 × 10^5^ cells per chip. Cell suspension was injected into the endothelial channel via the inlet port using a 200 μL pipette tip, with an empty tip placed at the outlet port to collect excess suspension. The cell suspension was perfused until it emerged from the outlet port tip, confirming channel filling. Devices were incubated at an inverted position for 3 hours to allow CD31^+^ cells to adhere to the porous membrane. Following incubation, all channels were carefully flushed with the corresponding cell type-specific medium with 1X Antibiotic Antimycotic (Cat. No. 15240096, Thermo Fisher Scientific). All media were pre-degassed for 24 hours at 37°C. Medium-filled 200 μL pipette tips were placed in all ports at the volumes specified in Table 1. All devices were transferred to a humidity chamber and incubated (37°C with 5 % CO₂) overnight.

**Table 1.** Tip volumes used for chip culture for each compartment.

| Channel | Medium | Inlet tip volume | Outlet tip volume |
| --- | --- | --- | --- |
| Neurofeeder channel | Maturation medium from STEMdiff™ Cerebral Organoid Kit | 30 µL | 40µL |
| Parenchymal compartment | Maturation medium from STEMdiff™ Cerebral Organoid Kit | 40 µL | - |
| Perivascular channel | 1:1 Astrocyte Medium and Pericyte Growth Medium 2 | 10 µL | 20 µL |
| Endothelial channel | Endothelial Cell Growth Medium MV2 | 50 µL | 50 µL |

On day 1, astrocytes and CD31^-^ pericyte-like cells were detached separately using 1X TrypLE, counted, and seeded at a density of 8 × 10^4^ cells per chip and 2 × 10^4^ cells per chip, respectively. Cells were introduced into the perivascular compartment following the same injection method used for CD31^+^ cells, with the exception that devices were maintained at an upright position during the 3-hour adhesion incubation. Following incubation, all channels were carefully perfused with fresh, degassed, cell-type specific medium with 1X Antibiotic Antimycotic following same volumes as on day 0.

On day 2, all channel media were exchanged for glucose-free medium, as described in Table 2. Channels were first washed with 20 µL of the corresponding glucose-free medium to remove residual glucose, after which fresh glucose-free medium was introduced via pipette tips placed in all ports except the endothelial channel inlet, which was connected to a peristaltic pump for continuous perfusion.

**Table 2.** Glucose-free media formulation.

| Channel | Medium |
| --- | --- |
| Neurofeeder channel and parenchymal compartment | 25 mL DMEM no glucose, 25 mL Neurobasal™-A Medium, no D-glucose no sodium pyruvate (A2477501, Thermo Fisher Scientific), 0,25 mL N2 supplement (17502048, Thermo Fisher Scientific), 12,5 µL Insulin (I9278, Sigma Aldrich), 1x GlutaMAX (35050061, Thermo Fisher Scientific), 0,25 mL MEM non/essential amino acids (M7145, Sigma Aldrich), 0,312 nM µL 2-mercaptoethanol (31350010, Thermo Fisher Scientific) , 0,5 mL B27 (17504044, Thermo Fisher, Scientific), 1X Antibiotic-Antimycotic |
| Perivascular channel | DMEM no glucose (11966025 , Thermo Fisher Scientific), 9% Pericyte Cell Growth Medium Supplement, 2% FBS, 1% AGS supplement from Astrocyte Medium, 1X Antibiotic-Antimycotic |
| Endothelial channel | DMEM no glucose (11966025 , Thermo Fisher Scientific), 5,42% Endothelial Cell Growth Medium Supplement, 1X Antibiotic-Antimycotic |

The perfusion circuit consisted of 5 mL Eppendorf tubes serving as inlet medium reservoirs, with their caps perforated at two points: one hole to accommodate an 18G metal connector and the second to serve as a pressure-release vent. A 5 cm length of transparent tubing (E3603, Tygon S3 E-lab 0.8mm 10/2.4 mm, Darwin) connected the reservoir to a gas-permeable peristaltic pump tubing segment (Tubing ISMATEC PharMed BPT 0.25mm ID Orange/Blue, MFLC95723-12, ISMATECH), which was in turn connected via an 18G metal connector to the endothelial channel inlet port. At the outlet port of the endothelial channel, an 18G connector was led into 11 cm of transparent tube directing effluent to the outlet collection reservoir of identical construction (Figure S2). The peristaltic pump flow was set to 1.716 µL/min. D-(+)-Glucose-^13^C_6_ (26707, Cayman Chemical Company) was added to the endothelial medium reservoirs to a final concentration of 5.3 mM to enable quantification of glucose transfer across compartments by targeted mass spectrometry.

On day 3, medium was replaced in all compartments and reservoirs. On day 4, after 24 hours of medium change, medium samples were collected from all compartments and stored at -20°C until further analysis. For RNA extraction, endothelial and perivascular channels were first washed with 50 µL cold DPBS^+^ to remove residual medium, after which 200 µL of lysis buffer (ReliaPrep™ RNA Miniprep Systems Kit, Z6011, Promega) was perfused through the endothelial channel inlet and collected at the outlet port into an empty 200 µL pipette tip. This process was repeated immediately for the perivascular channels, processing one chip at a time.

Cerebral organoids were carefully aspirated from the parenchymal compartment using a cut 1000 µL tip and transferred to a 1.5 mL Eppendorf tube containing 100 µL of cold DPBS^+^ to wash away residual Matrigel. Following aspiration of the DPBS^+^, 200 µL of lysis buffer was added and the cerebral organoid was mechanically lysed by repeated pipetting until no visible tissue remained. All lysate samples were stored at -80°C until further analysis. One chip per experimental round was fixed with 4% PFA for 45 minutes, washed with DPBS^+^, and stored at 4°C prior to performing immunocytochemistry.

#### 2.2.3 Control cell cultures

Two-dimensional control cultures were established in parallel with on-chip integration. CD31^+^ endothelial cells were seeded at 1.2x10^5^ cells per well in 24-well plates pre-coated with 0.084 mg/mL Matrigel (n=5). Astrocytes and CD31^-^ pericyte-like cells were similarly co-cultured in Matrigel-coated 24-well plates at the same seeding densities as used on-chip. Cerebral organoids were maintained on non-treated 6-well plates on an orbital shaker. Culture media for each cell type are specified in Table 1. Cell lysates were collected following the same procedure as for on-chip cultures, and one well per cell type was fixed with 4% PFA for immunocytochemistry.

### 2.3 Immunocytochemistry

Previously fixed samples were blocked and permeabilized using blocking buffer (10% normal goat serum, 0.1% Triton-X-100 in DPBS^+^ (with Mg^2+^ and Ca^2+^)) for 1 hour at room temperature. Primary antibodies (listed in Supplementary Table 3) were diluted in staining buffer (1% normal goat serum, 0.01% Triton-X-100 in DPBS^+^) and incubated overnight at 4°C. Samples were washed three times with DPBS^+^, after which the secondary antibodies, phalloidin and DAPI were diluted in staining buffer and incubated for 1 hour at room temperature. Stained samples were imaged using a laser-scanning microscope (Zeiss LSM 980, Carl Zeiss, Oberkochen, Germany) and Zen Blue software (v3.2, Carl Zeiss). Image post-processing was performed using ImageJ/Fiji (National Institutes of Health, Bethesda, United States). Brightness and contrast were adjusted for on-chip images based on signal intensities obtained from the corresponding 2D control samples to enable qualitative comparison of the cell morphology and distribution across conditions.

### 2.4 RNA isolation and RT-qPCR

Total RNA was extracted using ReliaPrep^TM^ RNA Miniprep Systems Kit (Cat. No. Z6010, Promega) according to the manufacturer’s instructions. RNA concentration and purity were assessed spectrophotometrically, and 1 µg of total RNA per sample was reverse-transcribed into cDNA using the High-Capacity RNA-to-cDNA™ Kit (Cat. No. 4387406; Thermo Fisher Scientific). Quantitative real-time PCR was performed using TaqMan™ Fast Advanced Master Mix (#4444963; Thermo Fisher Scientific) on a QuantStudio™ Real-Time PCR System (Thermo Fisher Scientific) or a CFX Real-Time PCR System (Bio-Rad), with each biological replicate analyzed in two or three technical replicates. RPL36AL and ACTB were used as endogenous reference genes. Gene expression was analyzed using the ΔΔCq method. In brief, (1) ΔCq, (2) ΔΔCq, and (3) fold change were calculated in Microsoft Excel using the following equations: 1) ΔCq = Cq gene of interest – Cq endogenous control; 2) ΔΔCq = ΔCq NEoC sample – ΔCq 2D culture sample; and 3) Fold change = 2-ΔΔCq. Gene expression results are represented as fold change (2^ΔΔCt) relative to the 2D culture. Fold change > 1 represents an upregulation of the gene, fold change <1 represents a downregulation of the gene, and fold change = 1 represents unchanged expression of the gene. Statistical analysis and data visualization were performed in OriginPro 2026, with p-values derived from using Linear Mixed Models (LMMs). Target genes and corresponding TaqMan™ assays are listed in Supplementary Table 5.

### 2.5 Targeted and untargeted mass spectrometry

Medium samples collected from each compartment of the NEoC were analyzed by targeted mass spectrometry for quantification of D-(+)-Glucose-^13^C_6_ to assess cross-compartment glucose transfer. Untargeted mass spectrometry was subsequently performed on the same samples to characterize the compartment-specific secretome and identify differentially abundant metabolites and enriched metabolic pathways. Briefly, raw data were processed using a Python-based metabolomics pipeline to predict metabolite enrichment, network activity, and identifies significantly affected pathways from high-throughput metabolomics. Full details on sample preparation, instrument parameters, data acquisition, and statistical analysis are provided in the Supplementary information.

## Supporting information

Supplementary Informattion

## Credit authorship contribution statement

JR conceived the NVU platform. MY, LB, and TW differentiated and characterized the CD31+ and CD31- cells. AC differentiated, cultured, and characterized COG. AC, LM, JV, MY performed biological experiments and immunofluorescence analysis. JV fabricated the microfluidic devices. LL, AC, LM performed qPCR analysis. JC performed computational simulations. JJ performed profilometry measurements. OF performed untargeted mass spectrometry analysis. KK performed targeted and untargeted mass spectrometry. SY, AWr, AWe contributed with the biological material. JR and AH conceived the study and the experimental design. AC, LM, JR, and AH wrote the manuscript.

## Acknowledgements

JR acknowledges funding by the European Union (HORIZON-MSCA-2022-PF-01, GA-101109010-NEoC). AH acknowledges funding by InfraPlus (Grand agreement 101131669).

## Data availability

The datasets generated for this study are available on request to the corresponding author.

