## Supplementary Informattion for "NeuroEnergetics-on-Chip: a novel 3-compartment microfluidic platform to study metabolic interactions between brain parenchyma and cerebral vasculature"

**Materials and methods**

**Supplementary Table 1.**Overview of key dimensions, areas, and volumes of the NEoC microfluidic platform.

| **Overall microfluidic platform dimensions** | | |
| --- | --- | --- |
| chip footprint | width [mm] | 21 |
|  | depth [mm] | 21 |
|  | area [mm^2^] | 441 |
| chip height | height of PDMS/PET layers [mm] | 7.71 |
|  | height of chip incl. 0.16 mm cover glass [mm] | 7.87 |
| **compartment 1a – *neurofeeder compartment*** | | |
| channel height | height [mm] | 0.08 |
| intersection chamber with *brain parenchyma compartment* | height [mm] | 0.20 |
|  | area [mm ^2^] | 2.00 |
|  | volume [µL] | 1.10 |
| entire compartment | area [mm ^2^] | 13.91 |
|  | volume [µL] | 2.49 |
|  | volume incl. ports [µL] | 14.05 |
| **compartment 1b – *brain parenchyma compartment*** | | |
| microchannel/micropillar dimensions | height [mm] | 0.02 |
|  | pillar width [mm] | 0.07 |
|  | channel width [mm] | 0.02 |
|  | pillar-/channel depth [mm] | 0.09 |
|  | number of microchannels (on one side) | 24 |
|  | number of microchannels (on both sides) | 48 |
| punching margin dimensions | height [mm] | 0.15 |
|  | pillar width [mm] | 0.07 |
|  | channel width [mm] | 0.19 |
|  | pillar-/channel depth [mm] | 0.35 |
|  | number of punching margin pillars (on one side) | 7 |
|  | number of punching margin pillars (on both sides) | 14 |
| open well structure | height [mm] | 4 |
|  | diameter [mm] | 3 |
|  | area [mm ^2^] | 7.07 |
|  | volume [µL] | 28.27 |
| entire compartment | area [mm ^2^] | 7.91 |
|  | volume [µL] | 49.60 |
|  | area without open well structure [mm ^2^] | 0.84 |
|  | volume without open well structure [µL] | 0.12 |
| **compartment 2 – *endothelial compartment*** | | |
| channel height | height [mm] | 0.20 |
| intersection area with *perivascular compartment* | area [mm ^2^] (on one side) | 2.45 |
|  | area [mm^2^] (on both sides) | 4.90 |
| entire compartment | area [mm^2^] | 17.47 |
|  | volume [µL] | 4.21 |
|  | volume incl. ports [µL] | 15.97 |
| **membrane** | | |
| membrane height | height [mm] | 0.011 |
| pore dimensions | pore size [mm] | 0.001 |
|  | porosity [%] | 1.6 |
|  | pore alignment | parallel |
| **compartment 3 – *perivascular compartment*** | | |
| channel height | height [mm] | 0.08 |
| intersection area with *endothelial compartment* | area [mm ^2^] (on one side) | 2.45 |
|  | area [mm ^2^] (on both sides) | 4.90 |
| entire compartment | area [mm ^2^] (on one side) | 5.29 |
|  | volume [µL] (on one side) | 0.42 |
|  | volume incl. ports [µL] | 11.42 |
|  | area [mm ^2^] (on both sides) | 10.58 |
|  | volume [µL] (on both sides) | 2 |
|  | volume incl. ports [µL] | 23.84 |

**NEoC Chip fabrication**

**3D printing of micropatterned master molds:** Master molds for the individual device layers were fabricated using a BMF Boston Micro Fabrication microArch® S140 printer in combination with the HTL-20 Resin (BMF Boston Micro Fabrication). Microstructures were designed on top of a 1 mm-thick 3D-printed substrate to provide mechanical support. For generating exclusion molding master molds, a 1 mm thick, 0.35 mm high rim was designed around the structure to contain the thin layers of PDMS during casting.

AutoCAD designs were exported as .stl files and converted to 10 µm-sliced exposure files using the BMF Slicer Software (version 1.6.4), with the parameters specified in Supplementary Table 2. To achieve the required 21 x 21 mm device footprint (exceeding the area of a single projection zone), Multi-Exposure (Stitch) Mode was selected, with a stitching overlap of 5 pixels and an exposure intensity fade factor of 0.4 to minimize stitch artefacts. Printed structures were uniformly scaled to 100.5% to compensate for material bleed-out inherent to HTL-20 resin.

**Supplementary Table 2.** Parameter settings in BMF Slicer Software for converting 3D-.stl files to slices for resin exposure in the BMF microArch® S140.

| **parameter** | **setting** |
| --- | --- |
| uniform scale for HTL [%] | 100.5 |
| slice thickness [µm] | 10 |
| pixel size [µm] | 10 |
| image resolution (for stitching function) | 9400x5200 |
| height [mm] | 1 + 0.35 (for neurofeeder & endothelial compartments)  1 + 0.5 (for top layer) |

Printing velocities were set to (i) 10.0 mm/s (with an acceleration of 10.0 mm/s^2^) for the XY-direction, (ii) 2.0 mm/s (with an acceleration of 1.0 mm/s^2^) for the platform down (P-d) velocity and (iii) 1.0 mm/s (with an acceleration of 1.0 mm/s^2^) for the platform up (P-u) velocity. Specific printing parameters for each layer of resin exposure, including exposure time and exposure intensity, are listed in Supplementary Table 3. The printer was calibrated prior to each print run according to the manufacturer's guidelines.

**Supplementary Table 3**. Printing parameters for 3D printing of standard and exclusion molding micropatterned master molds. Image: number of pictures/slices to be projected, with n = total number of pictures minus 35 pictures; Exp-t: exposure time; Intst: exposure intensity; P-d: distance by which the platform stage descends after exposure of individual layer*; Dt1: dwell time 1 – wait time after the platform stage descends; P-u: distance by which the platform stage ascends before exposure of next individual layer*; Dt2: dwell time 2 - wait time after the platform stage ascends; Scra: scraper movement by layers (e.g., Scra = 3 means scraper moves every 3 layers).

| **image** | **Exp-t [s]** | **Intst** | **P-d [mm]** | **Dt1 [s]** | **P-u [mm]** | **Dt2 [s]** | **Scra** |
| --- | --- | --- | --- | --- | --- | --- | --- |
| 1 | 4.00 | 50 | 0 | 0.00 | 0.000 | 1800.00 | 0 |
| 4 | 1.50 | 45 | 4.000 | 2.00 | 3.990 | 1200.00 | 3 |
| 10 | 1.00 | 45 | 4.000 | 2.00 | 3.990 | 800.00 | 5 |
| 20 | 1.00 | 45 | 4.000 | 2.00 | 3.990 | 600.00 | 5 |
| n | 1.00 | 45 | 4.000 | 2.00 | 3.990 | 250.00 | 5 |

*Remark: difference between P-d and P-u yields layer thickness. E.g., if P-d is 4.000 mm, and P-u is 3.990 mm, the layer thickness is 10 µm.

Following printing, the printing platform was carefully removed from the resin vat and directly transferred to an isopropanol bath, where it was soaked for 2 minutes with gentle manual agitation. Master models were detached from the printing platform using a razor blade held at a shallow angle to avoid surface damage, transferred to a fresh bath of isopropanol and then UV-cured for 1 hour at 60 °C (Formlabs UV Cure Station, Formlabs).

**Replica molding of PDMS layers:** The PDMS layers of the NEoC were fabricated by replica molding of PDMS using the 3D-printed micropatterned master molds (Figure S1). The top layer, comprising the structures of the perivascular and brain parenchyma compartments, was produced by standard molding, yielding a PDMS slab of approximately 5 mm thickness with the respective microstructures imprinted on its lower surface. For the neurofeeder and endothelial compartment layers, which require structures open on both sides, an exclusion molding approach was employed (**Figure S1b**).

PDMS (Sylgard 184 Silicone Elastomer Kit, Dow Corning) was mixed in a 10:1 (w/w) ratio of pre-polymer to curing agent and poured onto the respective master molds. The mixture was degassed under vacuum to remove air bubbles, ensuring a bubble-free replica molding. A thermoplastic release liner (3M Scotchpak™ 1022 Release Liner Fluoropolymer Coated Polyester Film; ID 70000200280,3M) was placed onto the uncured PDMS and pressed flat using a glass slide, then secured with clips. Master molds with PDMS were subsequently cured at 65° overnight. Upon cooling, the cured replica moldings were carefully released from the molds and inspected under a microscope to remove any residual particles or PDMS debris.

**PET membrane functionalization:** Isoporous ipCELLCULTURE™ PET membranes with 1 μm pore size (it4ip) were cut to the required dimensions using a 30 W CO_2_ laser cutter (VLS2.30, Universal Laser System Inc.), ensuring high precision and reproducibility across membranes. Prior to functionalization, membranes were immersed in 70% isopropanol and sonicated for 10 minutes to remove particle contamination, then dried at room temperature. Membranes were subsequently activated by oxygen plasma treatment (60 W for 60 s; CUTE-1MP/R, Femto Science) to introduce reactive surface groups. For silane functionalization, membranes were immersed in an isopropanol solution containing 2% (v/v) Bis[3-(trimethoxysilyl)propyl]amine (VWR), in individual small petri dishes placed on a hotplate at 80 °C, for 20 min, introducing primary amine groups at the membrane surface to enable covalent bonding to the PDMS layers during assembly. Membranes were then dried in the oven at 65°C for 30 minutes at, immersed in 70% ethanol for 30 minutes, and dried under a stream of compressed air immediately before assembly.

**Assembly of the multi-layered NEoC microfluidic platform:** The NEoC was assembled in a sequential, layer-by-layer bonding process (**Figure S1c**). First, a clean coverglass was taped to a glass slide for handling support. The neurofeeder PDMS layer and the coverglass were activated by O_2_ plasma treatment (15 s, 50 W), immediately bonded, and clamped between two glass slides. The assembly was placed at 65 °C overnight to consolidate attachment.

On the following day, the plasma-activated endothelial compartment PDMS layer, was aligned manually on top of the plasma-activated neurofeeder layer. The bonded pair was clamped between glass slides and again cured at 65 °C overnight.

On the third day, the functionalized PET membrane was positioned on top of the endothelial compartment PDMS layer. The top PDMS layer, comprising the parenchymal and perivascular compartments, was then plasma-activated and carefully aligned by hand to the underlying layers, with gentle lateral pressure applied against the PDMS edges to prevent collapse of the lateral microchannels. A piece of release liner was placed on top of the assembled stack, followed by a laser-cut PMMA (polymethyl methacrylate)-frame applied with light pressure to distribute clamping force evenly across the device surface. Approximately 10 glass slides were placed on top as a deadweight, and the complete assembly was cured at 65°C overnight.

Following the final curing, each device was inspected by bright-field microscopy. devices exhibiting collapsed microchannels or insufficient inter-layer adhesion were discarded.

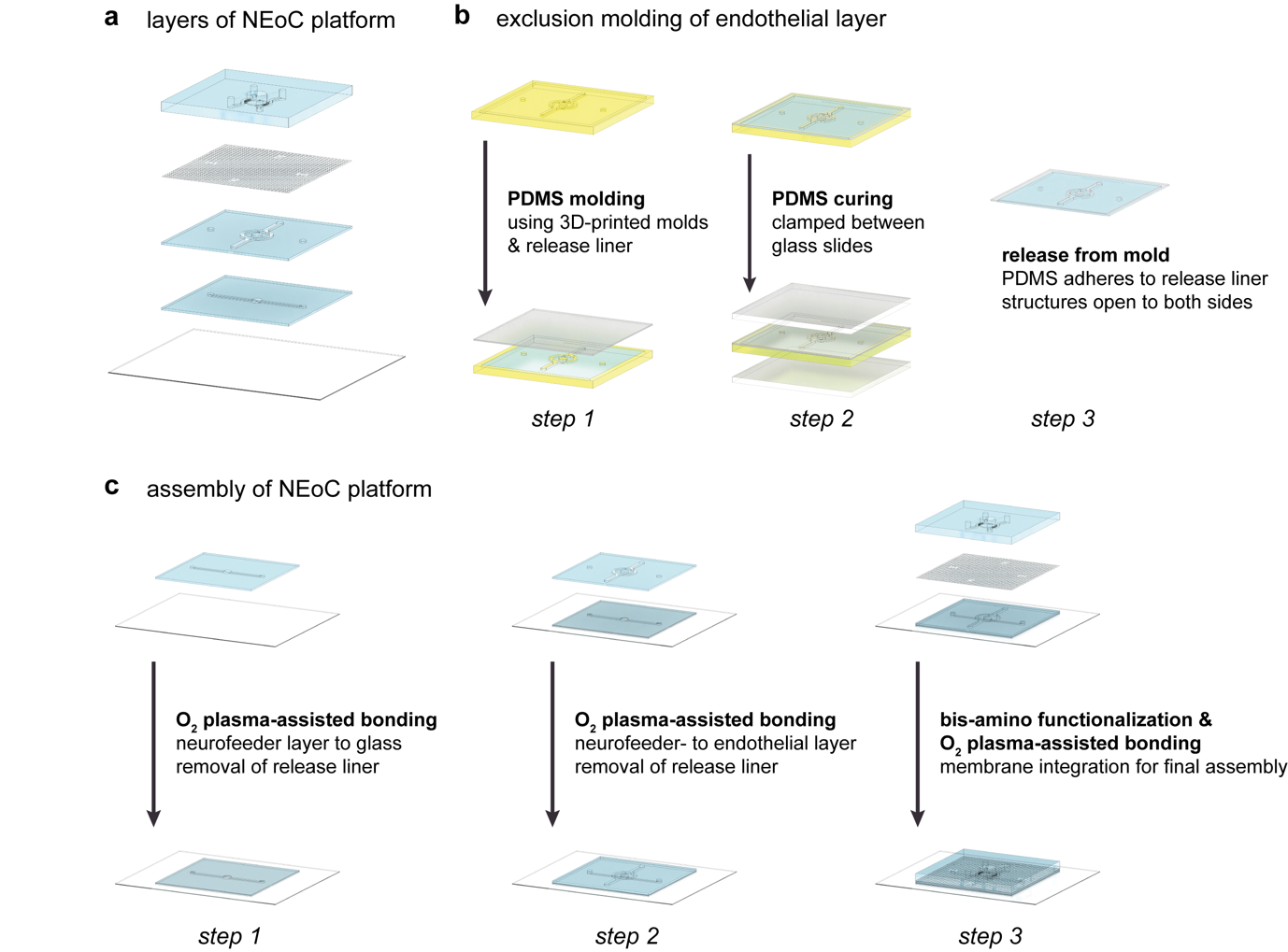

**Figure S1. Fabrication of the NEoC microfluidic platform**. **a.** Layered architecture of the microfluidic chip, from bottom to top: glass coverslip, neurofeeder PDMS layer (exclusion-molded), endothelial PDMS layer (exclusion-molded), isoporous PET membrane, and top PDMS layer containing the perivascular and brain parenchyma compartments. **b.** Exclusion molding process: uncured PDMS is poured into 3D-printed casting molds and covered with a release liner (step 1); the PDMS is cured beneath the release liner between two glass slides at 65 °C overnight (step 2); the cured PDMS is released from the 3D-printed mold while remaining adhered to the release liner, which is removed only during final assembly. **c.** Final assembly of the NEoC microfluidic platform. The neurofeeder layer is bonded to the glass coverslip via O₂ plasma-assisted bonding and clamped between two glass slides overnight at 65 °C (step 1). After removing the release liner from the neurofeeder layer, the endothelial layer is bonded on top via O₂ plasma-assisted bonding and similarly clamped overnight at 65 °C (step 2). The release liner is then removed from the endothelial layer. Immediately before final assembly, the PET membrane is functionalized with bis(3-(trimethoxysilyl)propyl)amine. Using O₂ plasma-assisted bonding, the functionalized membrane is sandwiched between the endothelial layer and the top layer and held under pressure overnight at 65 °C (step 3).

**Cell culture prior to on-chip culture**

**Human Induced Pluripotent Stem Cells** Human iPSC line Control 14 (C14 iPSC) used corresponds to a healthy female donor, provided by the iPS Core facility at Karolinska University Hospital. The use of human iPSCs was approved by the Swedish Ethical Review Authority (Etikprövningsmyndigheten, permit number: 2021-02050). Human iPSCs were cultured on surface-treated 6-well plates previously coated with 0,084 mg ml^-1^ Matrigel™ (#354230 Corning™ Matrigel™ Matrix) diluted in Dulbecco’s Modified Eagle Medium/Nutrient Mixture F-12 (DMEM-F12, Thermo Fisher, Waltham, MA, USA; Cat. No. 31331028), fed with mTeSR™ Plus medium (#100-0276, STEMCELL Technologies), and placed in a 37 °C, 5% CO2-supplemented incubator (#51033557, Heracell™ VIOS 160i CO2 Incubator, Thermo Fisher Scientific). A complete media change was performed the day after passage and every other day. Passaging was performed when confluency reached ~60-70%. Cells were treated with Versene solution (#15040033, Gibco, Thermo Fisher Scientific) for aggregate dissociation; after 4 min, the reaction was stopped with mTeSR™ Plus medium. Aggregates were detached by gently scratching the well plate surface with a 2 ml serological pipette. The cell suspension was mixed by gentle pipetting and seeded in a new, Matrigel-coated 6-well plate at the desired seeding ratio needed for further experiments.

**Endothelial cells and Pericytes** Endothelial-like cells (CD31+) and Pericyte-like cells (CD31-) were generated from the C14 iPSCs line following the protocol described by [38] with few modification. Generated cells were cryopreserved. After thawing, CD31⁺ and CD31⁻ cells were expanded in T75 flask previously coated with 0,2% gelatin (0423-SC, ScienCell) with Endothelial Cell Growth Medium MV2 (C-22022, Sigma-Aldrich, Merck) or Pericyte Growth Medium 2 (C-28041, Sigma-Aldrich, Merck) at 37 °C, 5% CO2-supplemented incubator, and used between passage 3-4. Cells were cultured until 80–90 % confluency before being seeded in the NEoC.

**Astrocytes** Astrocytes were generated from C14 iPSCs line. First neural epithelial stem cells (NES) were differentiated following the protocol from [39] and astrocytes were obtained following the protocol described by [40]. Generated cells were cryopreserved. Astrocytes were thawed and seeded in previously coated T25 flasks with 0,01% poly-L-ornithine (Sigma-Aldrich) and 10ug/ml laminin 2020 (Laminin from Engelbreth-HolmSwarm murine sarcoma basement membrane, Sigma-Aldrich) diluted in sterile DPBS+. Cells were maintained in Astrocyte Medium (#1801, ScienCell) at 37 °C with 5 % CO₂ in a humidified incubator. After 3 days of culture, the astrocytes were seeded into the microfluidic device.

**Cerebral Organoids** Cerebral organoids were differentiated from the C14 iPSCs line using the STEMdiff™ Cerebral Organoid Kit (#08570, STEMCELL Technologies) only modifying Rho-associated protein kinase inhibitor (ROCKi) concentration to 50 µM. In short, 90.000 cells/ml were seeded per well in a round-bottom 96-well plate. From day 1 to 4, Embryoid Body (EB) Formation Medium (Basal medium 1 and supplement A) was used. On day 5, EBs were transferred to a flat-bottom 48-well plate and cultivated in Induction Medium until day 7. On day 7, EBs were embedded in 15 µl of Matrigel (#354230, Corning® Matrigel® Growth Factor Reduced (GFR) Basement Membrane Matrix) and transferred to a non-treated 6-well plate with Expansion Medium. On day 10, the Expansion Medium was aspirated and replaced by the Maturation Medium, and the well plate was placed on an orbital shaker. Cerebral organoids were seeded on the microfluidic device at day 20.

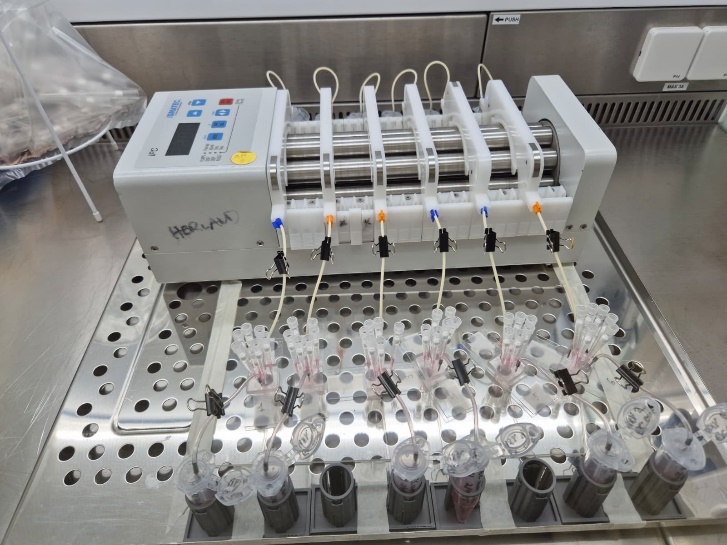

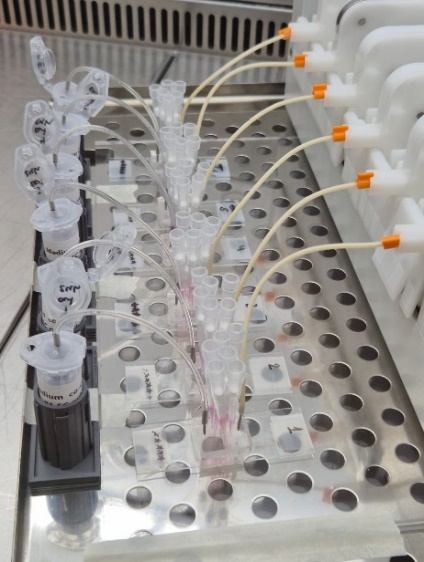

**Figure S2. Peristaltic pump connected to NEoC device.** Representative images.

**Supplementary Table 4.** Primary and secondary antibodies used for immunofluorescence procedures**.**

| **Primary antibody** | **Dilution** | **Supplier** | **Cat. No:** |
| --- | --- | --- | --- |
| Mouse anti-CD31 (PECAM1) monoclonal | 1:5 | Agilent Technologies | GA61061-2 |
| Rabbit anti-PDGFRβ monoclonal | 1:50 | Cell Signaling Technology | CST-3169S |
| Rabbit anti-vascular endothelial (VE)-cadherin monoclonal | 1:200 | Cell Signaling Technology | CST-2500S |
| Rabbit-Anti-beta III Tubulin | 1:200 | Abcam | ab18207 |
| Mouse anti-Glial fibrillary acidic protein | 1:500 | Sigma-Aldrich | AMAB91033 |
| Secondary antibody | Dilution | Supplier | Cat. No: |
| Goat anti-mouse IgG1 (γ1)  Alexa Fluor 488 polyclonal | 1:1000 | Sigma-Aldrich | SAB4600238 |
| Goat anti-rabbit IgG (H+L)  Alexa 633 | 1:1000 | Sigma-Aldrich | SAAB4600140 |
| Phalloidin Labeling Probes Alexa Fluor™ 555 | 1:400 | Thermo Fisher Scientific | A34055 |
| DAPI | 1:1000 | Invitrogen | D3571 |

**Supplementary Table 5.** Genes of interest for qPCR analysis**.**

| **Gene** | **Manufacturer** |
| --- | --- |
| *ACTB* | #4453320  Assay ID: Hs01060665_g1  Thermo Fisher Scientific |
| *RPL36AL* | #4448486 and #4331182  Assay ID: Hs00733231-m1  Thermo Fisher Scientific |
| *SLC2A1 (GLUT-1)* | #4331182  Assay ID: Hs00892681_m1  Thermo Fisher Scientific |
| *SLC2A3 (GLUT-3)* | #4453320  Assay ID: Hs00359840_m1  Thermo Fisher Scientific |
| *SLC16A1 (MCT1)* | #4331182  Assay ID: Hs01560299_m1  Thermo Fisher Scientific |
| *SLC16A7 (MCT2)* | #4331182  Assay ID: Hs00940851_m1  Thermo Fisher Scientific |
| *PECAM-1*  *(CD31)* | #4331182  Assay ID: Hs01065279_m1  Thermo Fisher Scientific |
| *CDH5 (VE*  *cadherin)* | #4453320  Assay ID: Hs00901463_m1  Thermo Fisher Scientific |
| *PDGFR-β* | #4453320  Assay ID: Hs01019589_m1  Thermo Fisher Scientific |
| *CSPG4 (NG2)* | #4448892  Assay ID: Hs05636647_s1  Thermo Fisher Scientific |
| *GFAP* | #4331182  Assay ID: Hs00909233_m1  Thermo Fisher Scientific |
| *S100B* | #4331182  Assay ID: Hs00902901_m1  Thermo Fisher Scientific |
| *DCX* | #4331182  Assay ID: Hs00167057_m1  Thermo Fisher Scientific |
| *TUBB3* | #4331182  Assay ID: Hs00964962_g1  Thermo Fisher Scientific |

**Mass spectrometry**

**Targeted mass spectrometry**

**Untargeted mass spectrometry** Untargeted metabolomics data were acquired in positive and negative ion mode by liquid chromatography-tandem mass spectrometry (LC-MS/MS). The dataset comprised 42 biological samples distributed across four experimental models, MonoCOG, BBB, NVU and NUVC9, each analyzed in two biological compartments, perivascular (peri) and parenchymal (COG), yielding eight experimental groups. A total of 568 annotated metabolite features were detected prior to filtering. Eight process blank samples (one per group) were included to support background signal subtraction. All preprocessing was implemented in Python (3.12.5) using pandas, NumPy, SciPy, and scikit-learn.

**SUPPLEMENTARY RESULTS**

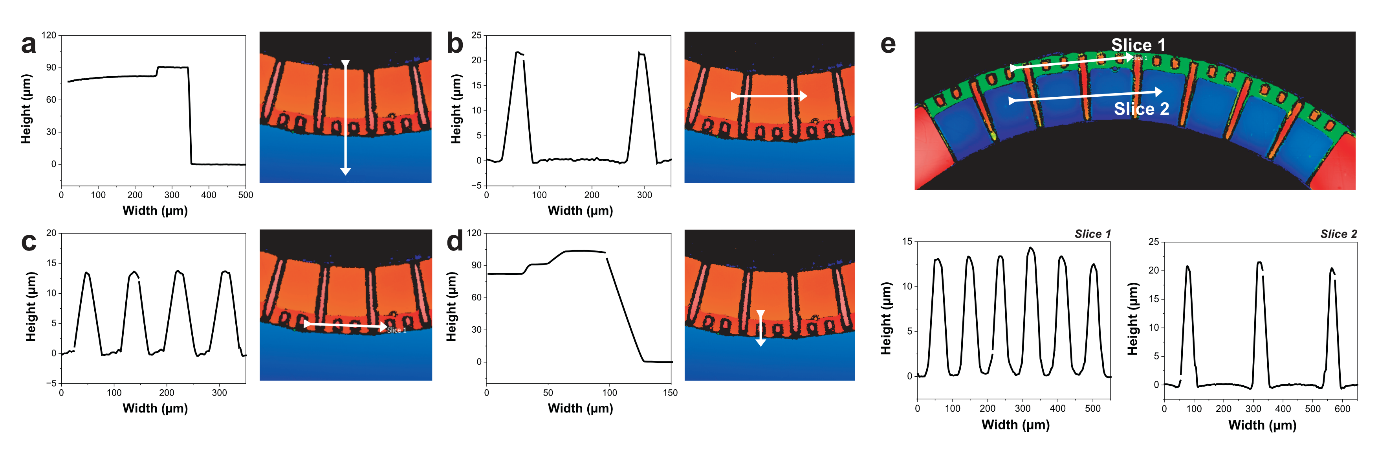

**Figure S3**. a-d. Dimensional measurements of the microchannels, micropillars, and punching margins (perivascular channel shown in blue) confirming high replication fidelity of the replica molding process. e.Cross-sectional profiles of the microchannels and micropillars.

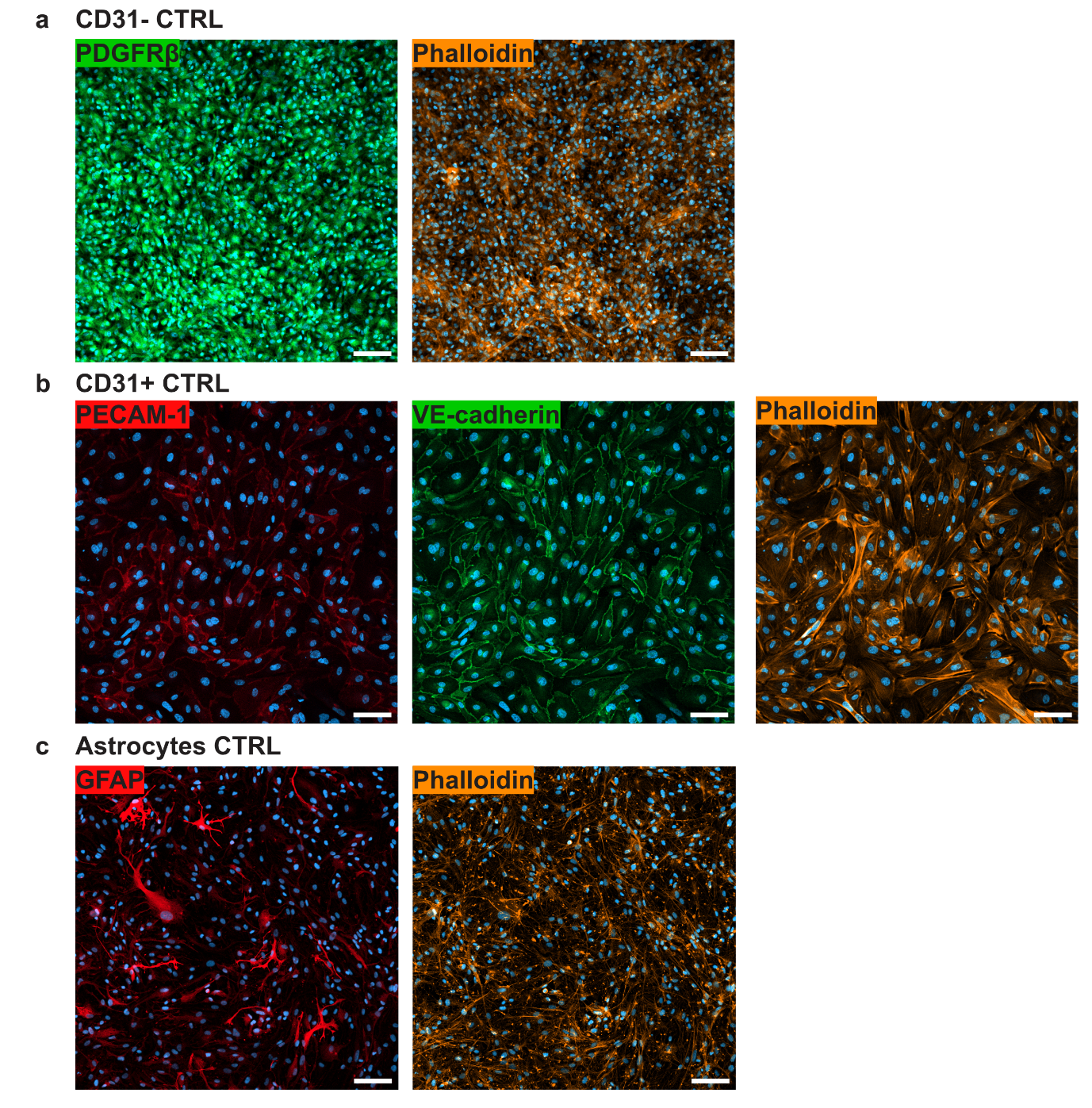

**Figure S4**. Immunofluorescence characterization of hiPSC-derived NVU cell types in 2D culture. **a.** CD31⁻ pericyte-like cells expressing PDGFRβ. **b.** CD31⁺ endothelial-like cells expressing PECAM-1 and VE-cadherin, visualizing adherens junctions. **c.** Astrocytes expressing GFAP. Scale bars equal 100 µm.

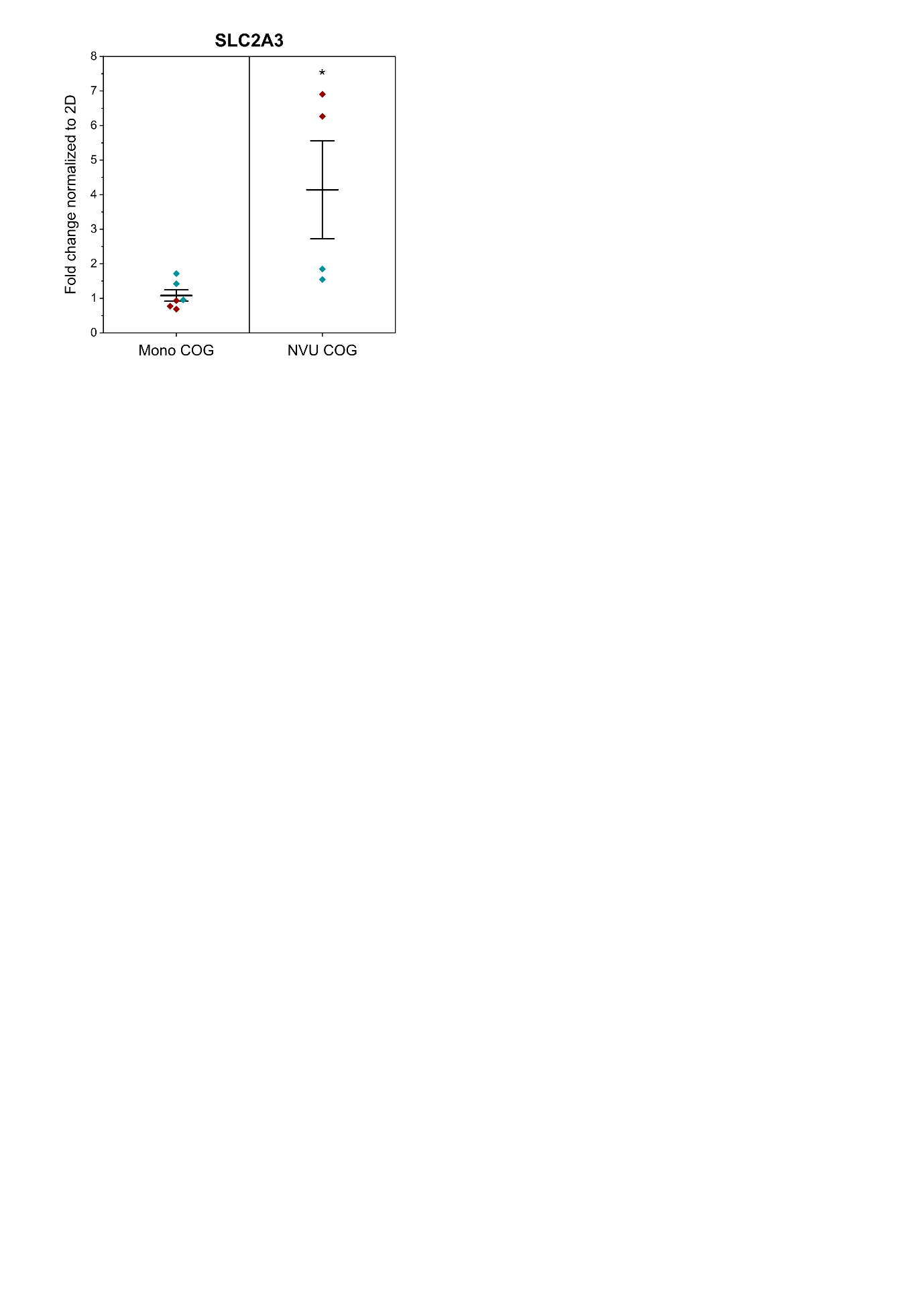

**Figure S5**. **GLUT3 expression in cerebral organoids across NEoC culture configurations.** *SLC2A3* mRNA expression measured in cerebral organoids under monoculture (*Mono COG*) and full *NVU* conditions. Data represented as fold change normalized to 2D culture controls. Whiskers indicate ± SE around the mean. P-values derived from linear mixed model analysis.

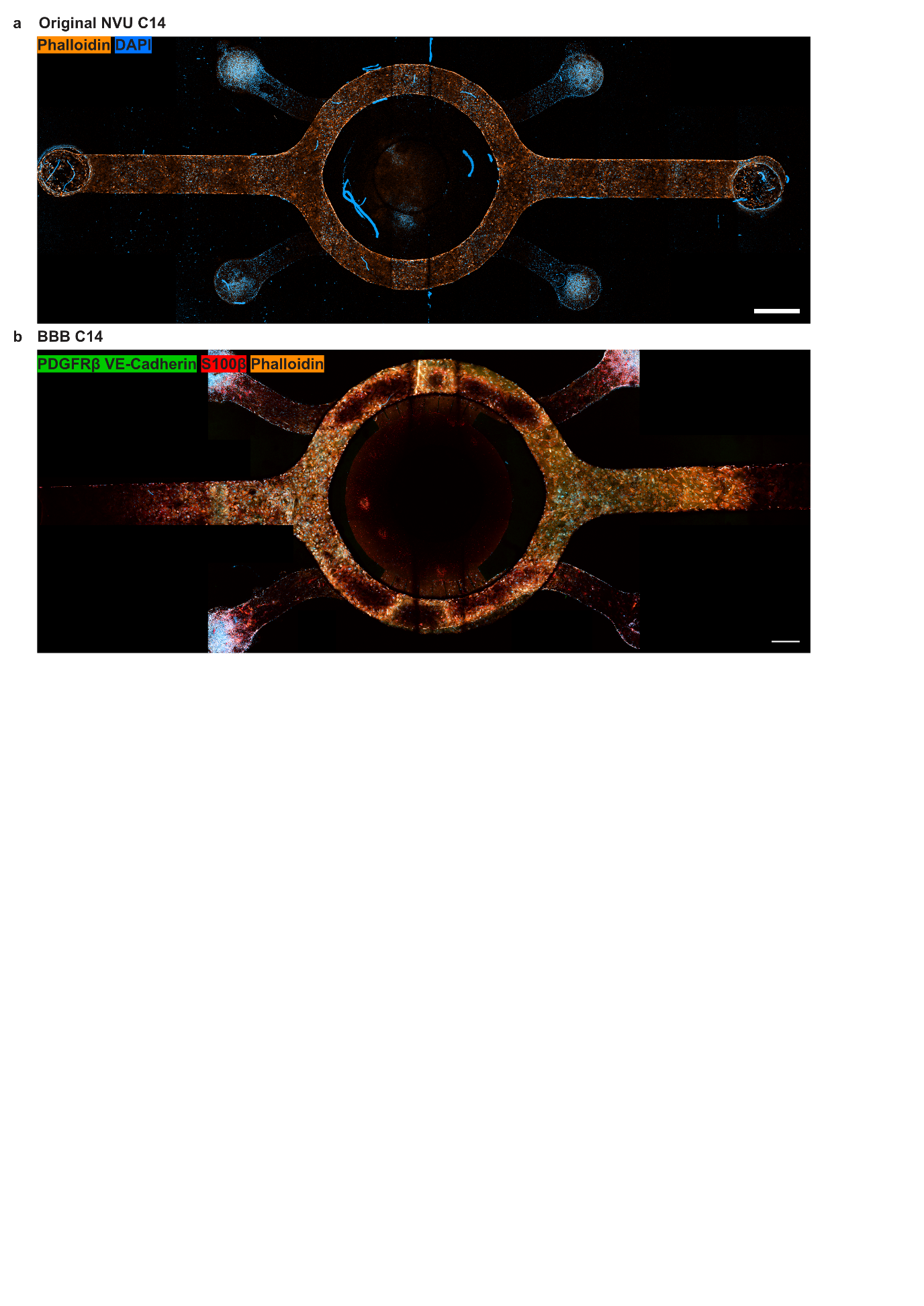

**Figure S6. Immunofluorescence imaging of hiPSC-derived NVU cell populations within the NEoC platform.** **a.** Full *NVU* condition stained with phalloidin (actin cytoskeleton, orange) and DAPI (nuclei, blue), showing the spatial distribution and morphology of all cell populations across compartments. **b.** *BBB* condition stained for VE-cadherin, PDGFRβ, and S100β, confirming the presence of endothelial-like, pericyte-like, and astrocytic populations within their respective compartments; cell-type identity of VE-cadherin⁺ and PDGFRβ⁺ cells is supported by compartmental location given the shared secondary antibody used for their detection. Scale bars equal 1000 µm.

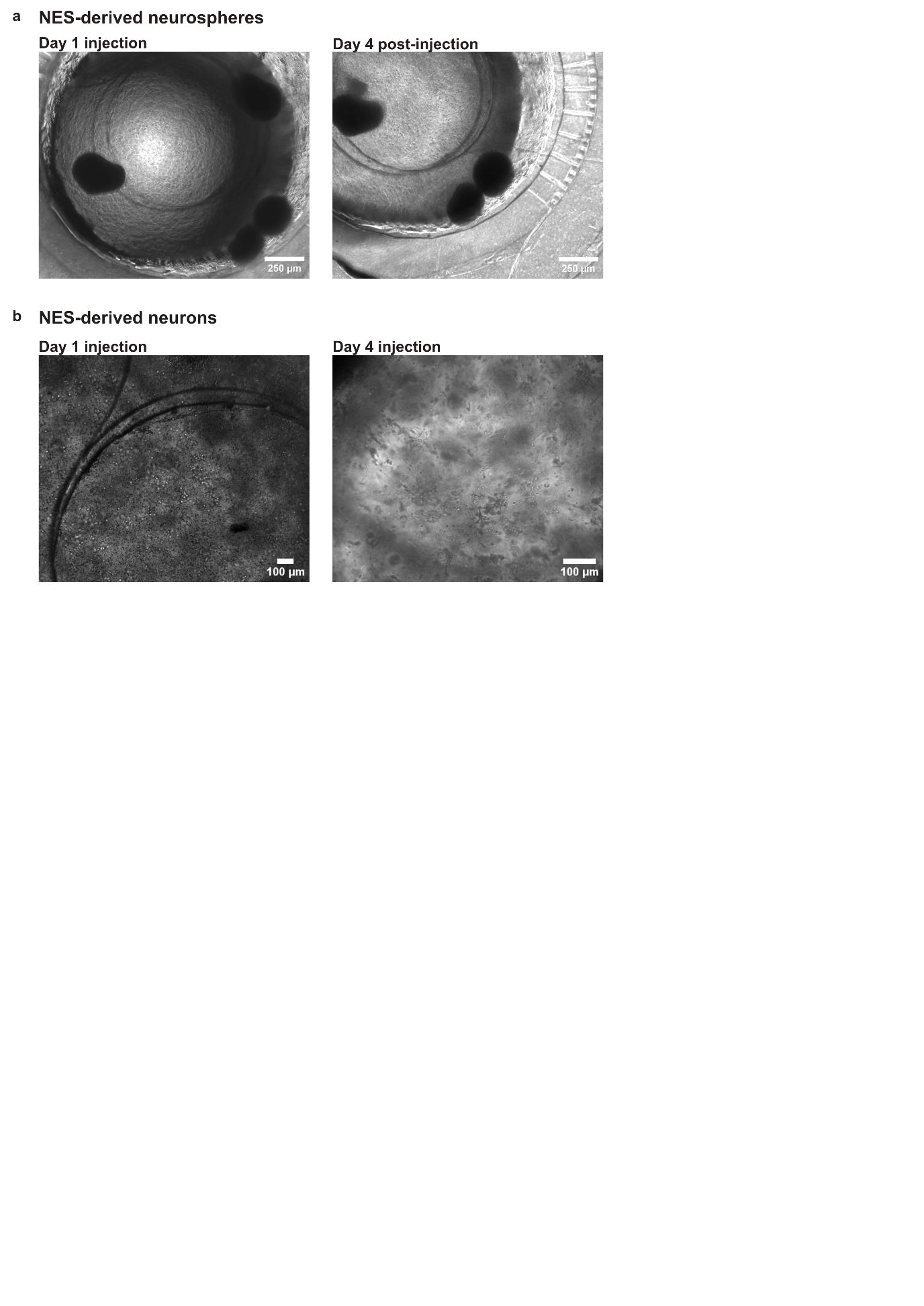

**Figure S7. Versatility of the parenchymal compartment: alternative 3D neural culture formats.** **a.** Representative image of neurospheres derived from neuroectodermal stem cells integrated within the parenchymal compartment. **b.** Representative image of dissociated neurons encapsulated in Matrigel hydrogel within the parenchymal compartment.

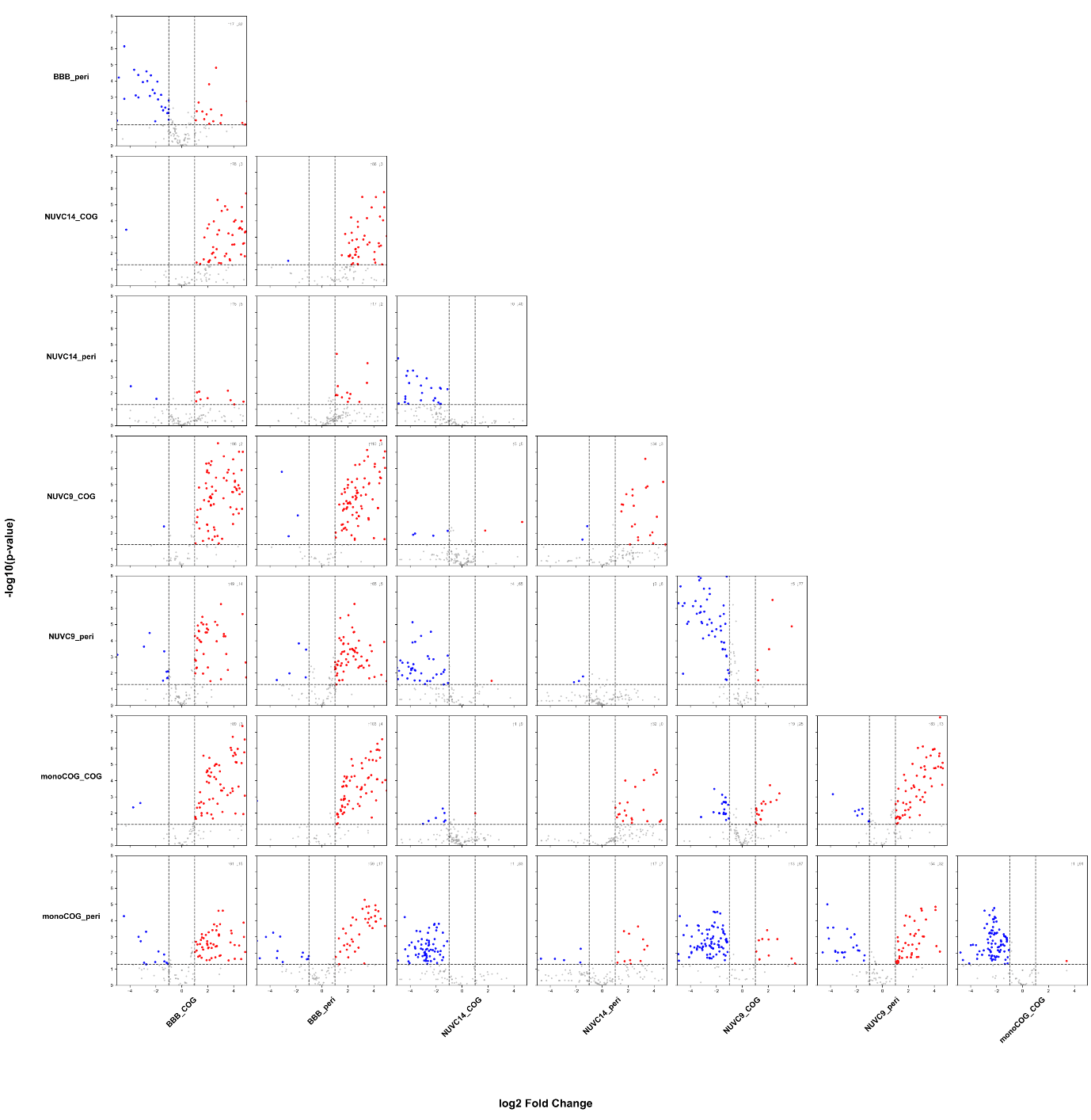

**Figure S8.** Pairwise differential abundance was assessed for all group combinations using Welch’s two-sample t-test applied to log_10_-transformed intensity values. Fold changes were computed as the difference in group means in log_10_ space converted to log_2_ scale: log_2_ FC = (mean_g2_ − mean_g1_ ) / log_10_ (2). Significance thresholds were set at p < 0.05 and |log 2 FC| > 1 (2-fold change). A multi-panel volcano plot and the key biological comparisons are shown. Significant hits for each comparison, including log 2 FC and p-values, are provided in **Supplementary Table 6**.
